# Major transcriptomic, epigenetic and metabolic changes underly the pluripotency continuum in rabbit preimplantation embryos

**DOI:** 10.1101/2021.10.06.463389

**Authors:** Wilhelm Bouchereau, Luc Jouneau, Catherine Archilla, Irène Aksoy, Anais Moulin, Nathalie Daniel, Nathalie Peynot, Sophie Calderari, Thierry Joly, Murielle Godet, Yan Jaszczyszyn, Marine Pratlong, Dany Severac, Pierre Savatier, Véronique Duranthon, Marielle Afanassieff, Nathalie Beaujean

**Affiliations:** Univ Lyon, Université Lyon 1, INSERM, Stem Cell and Brain Research Institute U1208, INRAE USC 1361, F-69500 Bron, France; Université Paris-Saclay, UVSQ, INRAE, BREED, 78350, Jouy-en-Josas, France; Ecole Nationale Vétérinaire d’Alfort, BREED, 94700, Maisons-Alfort, France; ISARA-Lyon, F-69007 Lyon, France; VetAgroSup, UPSP ICE, F-69280 Marcy l’Etoile, France; Université Paris-Saclay, CEA, CNRS, Institute for Integrative Biology of the Cell (I2BC), 91198, Gif-sur-Yvette, France; Univ. Montpellier, CNRS, INSERM, Montpellier France; Montpellier GenomiX, France Génomique, Montpellier, France

**Keywords:** Rabbit preimplantation embryo, pluripotency continuum, naive pluripotency, embryo transcriptome, single-cell RNAseq

## Abstract

Despite the growing interest in the rabbit model for developmental and stem cell biology, the characterization of embryos at the molecular level is still poorly documented. We conducted a transcriptome analysis of rabbit pre-implantation embryos from E2.7 (morula stage) to E6.6 (early primitive streak stage) using bulk and single-cell RNA-sequencing. In parallel, we studied oxidative phosphorylation and glycolysis and analysed active and repressive epigenetic modifications during blastocyst formation and expansion. We generated a transcriptomic, epigenetic, and metabolic map of the pluripotency continuum in rabbit preimplantation embryos and identified novel markers of naive pluripotency that might be instrumental for deriving naive pluripotent stem cell lines. Although the rabbit is evolutionarily closer to mice than to primates, we found that the transcriptome of rabbit epiblast cells shares common features with that of humans and non-human primates.

**Summary Statement:** Rabbit preimplantation embryos share characteristics with human and monkey embryos with respect to timing of early lineage segregation and expression of marker genes for naive and primed pluripotency.

## Introduction

In mammalian embryos, totipotent blastomeres become pluripotent after differentiation of the trophectoderm lineage during the morula to blastocyst transition, and form a seemingly coherent cluster of cells called the inner cell mass (ICM). When some ICM cells differentiate to the primitive endoderm (PE), pluripotency is confined to the epiblast (EPI), which will give rise to the embryo proper. Primordial germ cells (PGCs) are specified in the epiblast. thereafter Ultimately, the posterior part of the epiblast gives rise to the primitive streak (PS) (Blakeley et al., 2015; Chazaud and Yamanaka, 2016; Hayashi et al., 2007; Nakamura et al., 2016; Petropoulos et al., 2016; Stirparo et al., 2018). The developmental window that extends from morula differentiation to gastrulation lasts 2–3 days in mice, compared to 7–10 days in humans and non-human primates (Nakamura et al., 2021; Shahbazi, 2020). Within that window, pluripotent cells are thought to transit through three main pluripotency states known as the naive, formative and primed states (Savatier et al., 2017; Smith, 2017; Takahashi et al., 2017). In mice, the ICM and epiblast cells of blastocysts, as well as the embryonic stem (ES) cell lines that are derived from them, are in the naive state. Naive pluripotency is associated with the expression of transcription factors Klf2, Klf4, Gbx2, Tfcp2l1, Esrrb, Tbx3, and Sall4 (Dunn et al., 2014), low DNA methylation (Leitch et al., 2013; Leitch et al., 2016), high mitochondrial respiration (Carbognin et al., 2016; Sone et al., 2017), and enrichment of active histone modifications at promoter regions of developmental genes (Hayashi et al., 2008). Formative pluripotency characterizes the epiblast cells of peri-implantation embryos, as well as the corresponding pluripotent stem cell lines, called formative stem (FS) cells (Kinoshita et al., 2020; Smith, 2017). The formative state is associated with the expression of transcription factors Etv5, Rbpj, Tcf3, and Otx2, and intermediate DNA methylation (Kalkan et al., 2019; Kinoshita et al., 2020). Finally, the epiblast cells of early to late gastrula-stage embryos and their *in vitro* counterparts, epiblast stem cells (EpiSCs), are in the primed pluripotent state (Brons et al., 2007; Tesar et al., 2007). The primed state is associated with expression of Sox3 and Oct6 transcriptional regulators (Corsinotti et al., 2017), high DNA methylation (Habibi et al., 2013; Hackett et al., 2013), low mitochondrial respiration (Zhou et al., 2012), and enrichment of repressive histone modifications at promoter regions of developmental genes (Hayashi et al., 2008; Smith, 2017). These three successive pluripotent states characterize the pluripotency continuum.

The characterization of the mammalian embryo transcriptome in a wide range of species is essential for unravelling the molecular mechanisms involved in pluripotency and understanding the adaptation of embryos in maintaining pluripotency across different developmental strategies. It is also important for the development of chemically-defined culture media, which can aid in the preservation and maintenance of naive, formative, and primed pluripotency in embryo-derived stem cell lines. Single-cell transcriptomic data from preimplantation embryos is available in a large variety of species, including mice (Argelaguet et al., 2019; Mohammed et al., 2017; Tang et al., 2010), pigs (Kong et al., 2020), cattle (Zhao et al., 2016), marmosets (Boroviak et al., 2018), rhesus and cynomolgus macaques (Liu et al., 2018; Nakamura et al., 2016), and humans (Blakeley et al., 2015; Petropoulos et al., 2016). Transcriptome data at pre- and mid-gastrula stages is only available in mice (Peng et al., 2016; Wen et al., 2017) and cynomolgus macaques (Nakamura et al., 2016). It is indeed difficult to study this developmental period, as embryos have already implanted by the gastrula stage in many species. In mice, pigs, cattle, humans, and non-human primates, gastrulation starts two to eight days after implantation, when embryos are already deeply buried in the uterine wall, making the formative- and primed-pluripotency states difficult to study (Nakamura et al., 2021; Shahbazi, 2020). In contrast, rabbit embryos do not implant until the end of day seven of development, after the onset of gastrulation (Fischer et al., 2012; Puschel et al., 2010). This strategy of lagomorph development allows easier access to a larger window of development compared to rodents and primates. However, the transcriptome of the rabbit embryo is poorly documented (Leandri et al., 2009; Schmaltz-Panneau et al., 2014). For this reason, we conducted a transcriptome analysis of rabbit pre-implantation embryos between E2.7 (morula stage) and E6.6 (early primitive streak stage) using two complementary techniques: bulk RNA-sequencing (Illumina NextSeq) and single-cell RNAseq (10x Genomics)-to study specific gene expression. In addition, we studied oxidative phosphorylation and glycolysis, and analysed epigenetic modifications. With this unique combination of tools and cellular processes, we explored in detail the major alterations that occur in pluripotent cells *in vivo*, along the pluripotency continuum of rabbit embryos.

## Results

### Rabbit preimplantation embryos show both similarities and differences with mice and macaques in lineage marker expression

To obtain a first transcriptional map of the rabbit preimplantation embryo development, we first performed high depth bulk RNA sequencing of micro-dissected embryos between embryonic day E2.7 (morula) and E6.6 (expanded blastocyst at the early primitive streak-stage) (**Figs 1A, S1A**). The 51 samples –namely morula, trophectoderm (TE), inner cell mass (ICM), epiblast at E6.0 (EPI), anterior epiblast at E6.3/6.6 (EPIant), primitive endoderm (PE) resulting from these dissections separated according to developmental time and lineage when projected onto the first two components of principal component analysis (PCA) (**Fig. 1B**). We then investigated the expression of some known lineage markers of mouse, human, and cynomolgus macaque embryos, and represented the results on a heatmap (**Fig. 1C**). As expected, the trophectoderm (TE; E3.5 to E6.6) and inner cell mass (ICM; E3.5 and E4.0) were enriched in trophectoderm (e.g. *TFAP2C, KRT18*) and pluripotency (e.g. *DPPA5, ESRRB, POU5F1 (OCT4), SOX2*) transcripts, respectively. Most of the pluripotency genes had lower transcript levels in epiblast samples (EPI at E6.0, E6.3, and E6.6, thereafter called EPI_6.0, EPIant_6.3 and EPIant_6.6, respectively), which showed higher expression of late epiblast markers (e.g. *OTX2*). Finally, the primitive endoderm samples (PE at E6.0, thereafter called PE_6.0) were enriched in PE-specifi*c* transcripts such as *SOX17* and *GATA6*. Both TE markers (*GATA3* and *TFAP2C*) and ICM markers (such as *DDPA5, SOX15, KLF4, STAT3, KLF17, ESRRB*) were enriched in the E2.7 morula samples, suggesting early commitment of blastomeres either to ICM or TE cells. PE markers including *PDGFRA, GATA6, HNF1B, FOXA2, RSPO3*, and *SOX17* were detected in the E3.5 and E4.0 ICM samples suggesting that the PE differentiation begins as early at E3.5. Note that *FBXO15* and *TDGF1/CRIPTO*, two markers of naive pluripotency in mice (Boroviak et al., 2015; Mohammed et al., 2017), were up-regulated in rabbit EPI samples. Conversely the TE markers *PLAC8* or *SLC7A4* expressed in humans (Blakeley et al., 2015) were not found in rabbit TE. On the other hand, the primate-specific EPI markers *ZIC3* and *FGF2* (Nakamura et al., 2016), as well as the mouse-specific EPI markers *FGF5* and *SOX4* (Mohammed et al., 2017), were all expressed in rabbit EPI samples. Overall, these observations suggest that rabbit gene expression patterns share characteristics with both rodents and primates, though there are some notable differences.

**Figure 1:**
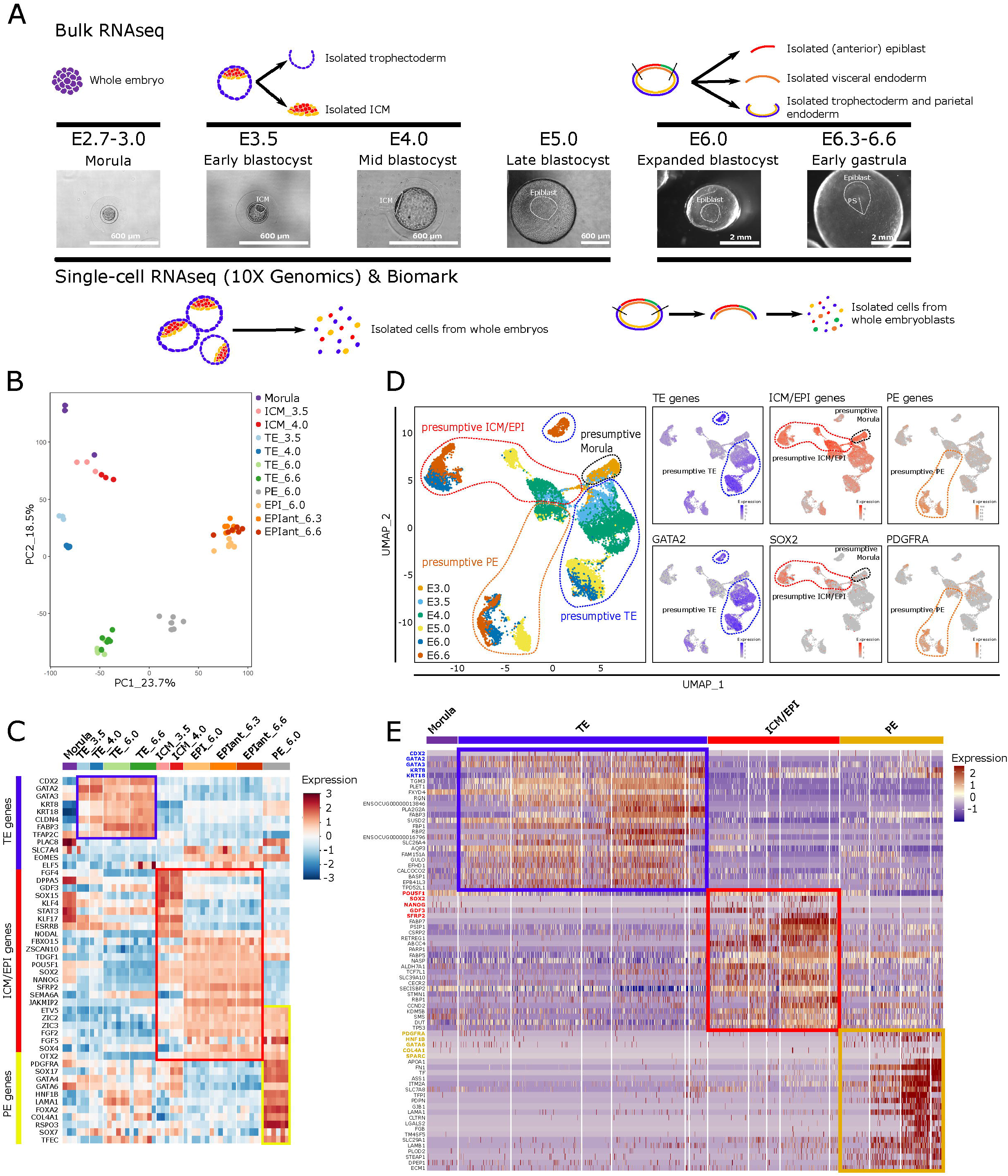
Transcriptional map of rabbit preimplantation embryo. (**A**) Experimental design with sample description. E2.7 to E6.6 indicate days of development after coitum (or insemination); ICM: inner cell mass; PS: primitive streak. **(B**) Graphical representation of the two first principal components of PCA for the 51 sample transcriptomes generated using bulk RNAseq data. (**C**) Heatmap representation of the expression level of 48 genes selected based on their known expression in early embryonic lineages of rodents and primates (bulk RNAseq data). (**D**) Two-dimensional UMAP representations of 13,942 single-cell transcriptomes (left panel) and showing marker gene expression (right panels). TE genes include *CDX2*, *CLDN4*, *FABP3, GATA2, KRT8, KRT18* and *TFAP2C*; ICM/EPI genes include *FGF4, GDF3, NANOG, SEMA6A*, *SFRP2*, *SOX2* and *SOX15*; PE genes include *COL4A1*, *FOXA2*, *GATA4*, *GATA6*, *HNF1B, LAMA1, PDGFRA* and *SOX17*. E3.0–E6.6 indicate the embryonic day (10x Genomics data). Dashed lines indicate presumptive lineage identities. (**E**) Heatmap representation grouping 20 gene markers (from B and C, highlighted in blue, red and yellow) with the 20 most differentially expressed genes in TE *vs*. all the other cell types (all), ICM and EPI *vs*. all, and PE *vs*. all (from D).

We next used the 10x Genomics technology to refine the transcriptional map of the rabbit preimplantation embryo development at a single-cell resolution (**Fig. 1A**). This technology is best suited for analysing large numbers of cells, although at a lower sequencing depth than bulk RNAseq (Nowotschin et al., 2019). Note that this study included additional E5.0 embryos that could not be included in the bulk RNAseq study. After quality control, we retained 13,942 single-cell transcriptomes from 529 embryos, ranging from E3.0 (morula) to E6.6 (early primitive streak-stage) (**Fig. S1B-D**). We applied uniform manifold approximation and projection (UMAP) dimension reduction and found that cells separated according to developmental stage when projected onto the first two components (**Fig. 1D**, **Table S1**). The six embryonic day clusters separated further into smaller clusters as follows: E3.0 cells (morula stage) formed one cluster. Cells harvested from the E3.5 early blastocysts formed two distinct clusters. The first one was enriched in ICM/EPI genes (e.g. *SOX2*) while the other cluster was enriched in TE genes (e.g. *GATA2*). At E4.0, cells separated into three clusters: the presumptive ICM/EPI and TE clusters, and a third cluster corresponding to presumptive PE represented by *PDGFRA* expression. At E4.0, the presumptive EPI and PE clusters were still close to each other. In contrast, at E5.0 the three lineages formed well-separated clusters. Based on these observations, we were able to delineate with certainty the morula, ICM, EPI, TE and PE clusters on the UMAPs.

To have a more global view of differential gene expression between the main cell clusters, a heatmap was generated using five landmark markers (**Fig. 1E** genes in colour fonts), with the 20 most overexpressed genes in TE *vs*. all, ICM/EPI *vs*. all, and PE *vs*. all comparisons (**Fig. 1E, Table S1**). Most of the genes represented on the heatmap had similar expression profiles to those described in mouse, cynomolgus macaque, and human (Guo et al., 2010; Mohammed et al., 2017; Nakamura et al., 2016; Stirparo et al., 2018). However, two genes showed unexpected expression profile. Namely, *SUSD2*, a marker of ICM in primates (Bredenkamp et al., 2019) is highly expressed in rabbit TE, and *CCND2*, a marker of the primitive streak in mice (Wianny et al., 1998) is already expressed in E5.0 EPI cells in rabbits. Finally, we integrated our dataset with published single-cell data from mouse and cynomolgus macaque (Nakamura et al., 2016) and thus confirmed the identity of the rabbit clusters (**Fig. S2**). Taken together, these results indicate that rabbit preimplantation embryos typically express the same lineage markers as mouse and cynomolgus macaque embryos, but there are also noticeable differences.

### EPI, PE, and TE lineages are established at E5.0 in rabbit embryos

To obtain a dynamic view of lineage specification, we performed pseudo-time analyses of the single-cell RNAseq dataset based on the 1,000 most variable genes. A first pseudo-time analysis was performed with the morula, ICM/EPI_3.5/5.0 and TE 3.5-5.0 cells (**Fig. 2A**). The morula cells (E3.0) split into two main branches at E3.5, leading to either EPI or TE cells at E4.0 (**Fig. 2B)**. At E5.0, EPI and TE cells formed two distinct clusters at the tip of their respective branches. We also identified two minor clusters located between the main clusters **(Fig. 2A)** and connected to the TE branch **(Fig. 2B)**; we qualified those cells as “TE intermediates”. A heatmap (**Fig. 2C**) was generated grouping 10 landmark genes with the 20 most overexpressed genes in the ICM/EPI *vs*. all and TE *vs*. all (**Table S1**). Up-regulated genes in TE *vs*. EPI lineage included *TFAP2C, GATA2, AQP3* and *CDX2* (**Fig. 2C**). Immunostaining of rabbit embryos confirmed ICM/EPI-specific expression of SOX2 at E3.5 and E4.0, and TE-enriched expression of TFAP2C at E4.0 (**Fig. 2D**).

**Figure 2:**
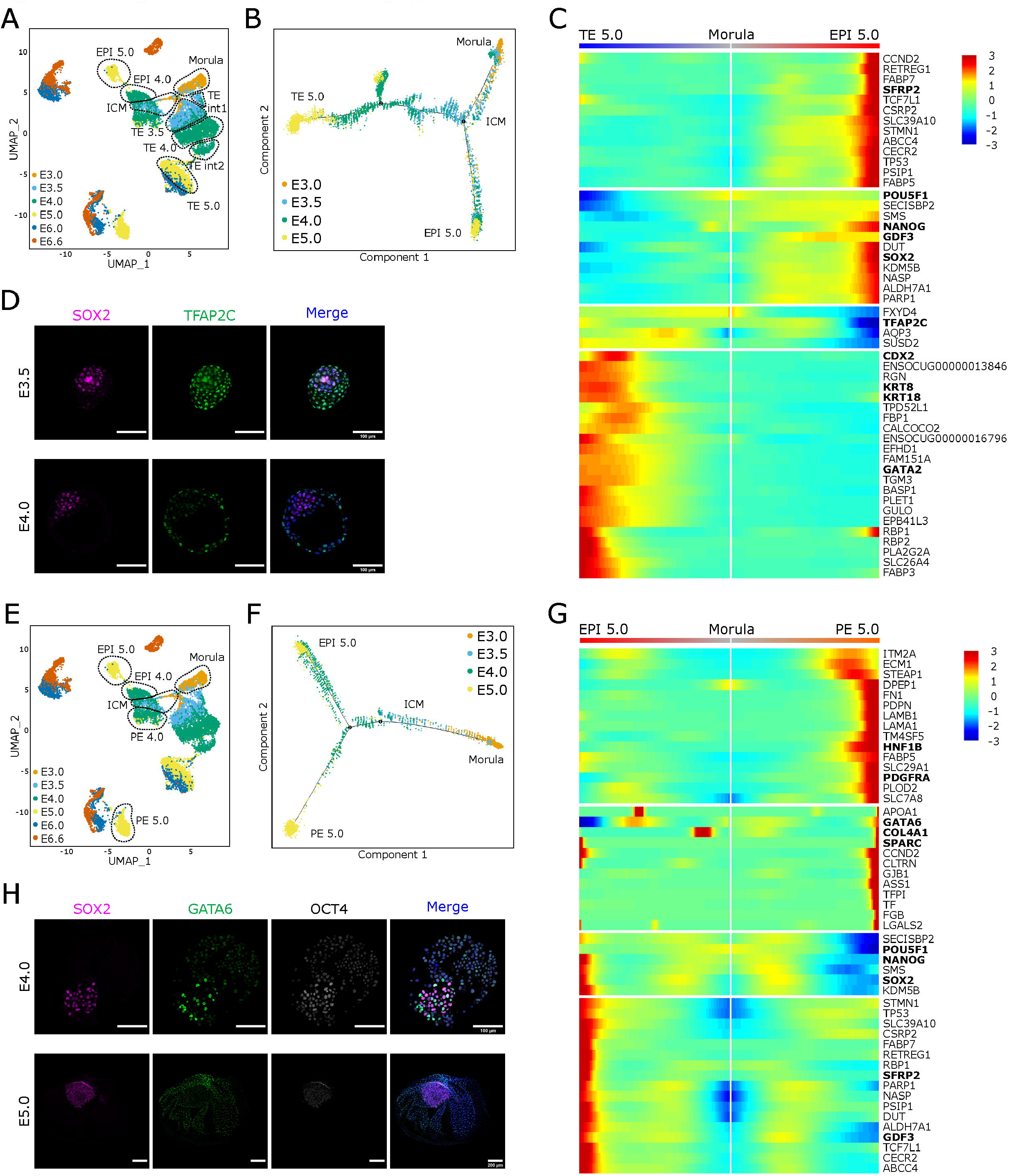
Timing of ICM/TE and EPI/PE segregation. (**A**) Two-dimensional UMAP representation of the 13,942 single-cell transcriptomes showing cells selected for the first pseudo-time analysis (dashed lines). **(B**) Pseudo-time representation of selected cells (shown in A), organized by embryonic day. (**C**) Heatmap representation of 10 landmark genes for the ICM/EPI (*POU5F1*, *NANOG*, *SOX2*, *GDF3*, *SFRP2*) and TE lineage (*CDX2*, *GATA2*, *TFAP2C*, *KRT8* and *KRT18*), and 20 most differentially expressed genes in the ICM/EPI and TE *vs*. all. (**D**) Immunofluorescence analysis of SOX2 and TFAP2C in E3.5 (n = 7) and E4.0 (n = 4) blastocysts. Merge pictures include DAPI staining. Scale bars: 100 μm. (**E**) Two-dimensional UMAP representation of the 13,942 single-cell transcriptomes showing cells selected for the second pseudo-time analysis. (**F**) Pseudo-time representation of selected cells (shown in E) organized by embryonic day. (**G**) Heatmap representation of 10 landmark genes for the ICM/EPI (*POU5F1*, *NANOG*, *SOX2*, *GDF3*, *SFRP2*) and PE lineage (*PDGFRA*, *HNF1B*, *GATA6*, *COL4A1* and *SPARC*), and 20 most differentially expressed genes in the ICM/EPI and PE *vs*. all. (**H**) Immunofluorescence analysis of SOX2, OCT4, and GATA6 at E4.0 (n = 4) and E5.0 (n = 4). Merge pictures include DAPI staining. Scale bars: 100 μm at E4.0 and 200 μm at E5.0.

In the second pseudo-time analysis, we used the same dataset except that the TE cells were replaced by the PE 4.0-5.0 cells (**Fig. 2E**). This analysis revealed a clear split of embryonic cells into two lineages, EPI and PE respectively, from E4.0 onwards (**Fig. 2F)**. At E5.0, EPI and PE cells formed two distinct clusters at the tip of their respective branches. A heatmap (**Fig. 2G)** was generated grouping 10 landmark genes with the 20 most overexpressed genes in the ICM/EPI *vs*. all and PE *vs*. all (identified in **Fig. 1E, Table S1**). Up-regulated genes in PE *vs*. EPI lineage included *GATA6, LAMA1, SPARC* and *PDGFRA* (**Fig. 2G**). Immunostaining of rabbit embryos confirmed PE-enrichment of GATA6 *vs* EPI-specific localization of SOX2 and EPI-enrichment of OCT4 at E4.0 and E5.0 (**Fig. 2H**). Together, these results confirm the establishment of the three main lineages, EPI, PE, and TE at E5.0 in rabbit embryos.

### Specification of visceral/parietal endoderm and anterior/posterior EPI takes place after E6.0

We sought to characterize the segregation of endodermal cells into visceral and parietal endoderm (VisE and ParE, respectively) using single-cell RNAseq data from PE cells (**Fig. 3A**). At E4.0 and E5.0, expression of the PE-specific gene *OTX2* gene was quite homogeneous among endodermal cells. However, at E6.0 and E6.6, endodermal cells formed two closely related clusters and *OTX2* expression was then restricted to one of these two subgroups (**Fig. 3A**). To characterize these *OTX2*-positive cells, we performed immunofluorescence analysis of E6.0 embryos. Strong OTX2 signal was only observed in SOX2-negative VisE cells underlying the layer of SOX2/OTX2 double-positive EPI cells (**Fig. 3B**). In marmosets, *OTX2* expression is also suppressed in the ParE and only detected in the VisE (Boroviak et al., 2018). These results suggest that segregation of visceral and parietal endoderm takes place at E6.0 in rabbit embryos, i.e. before embryo implantation.

**Figure 3:**
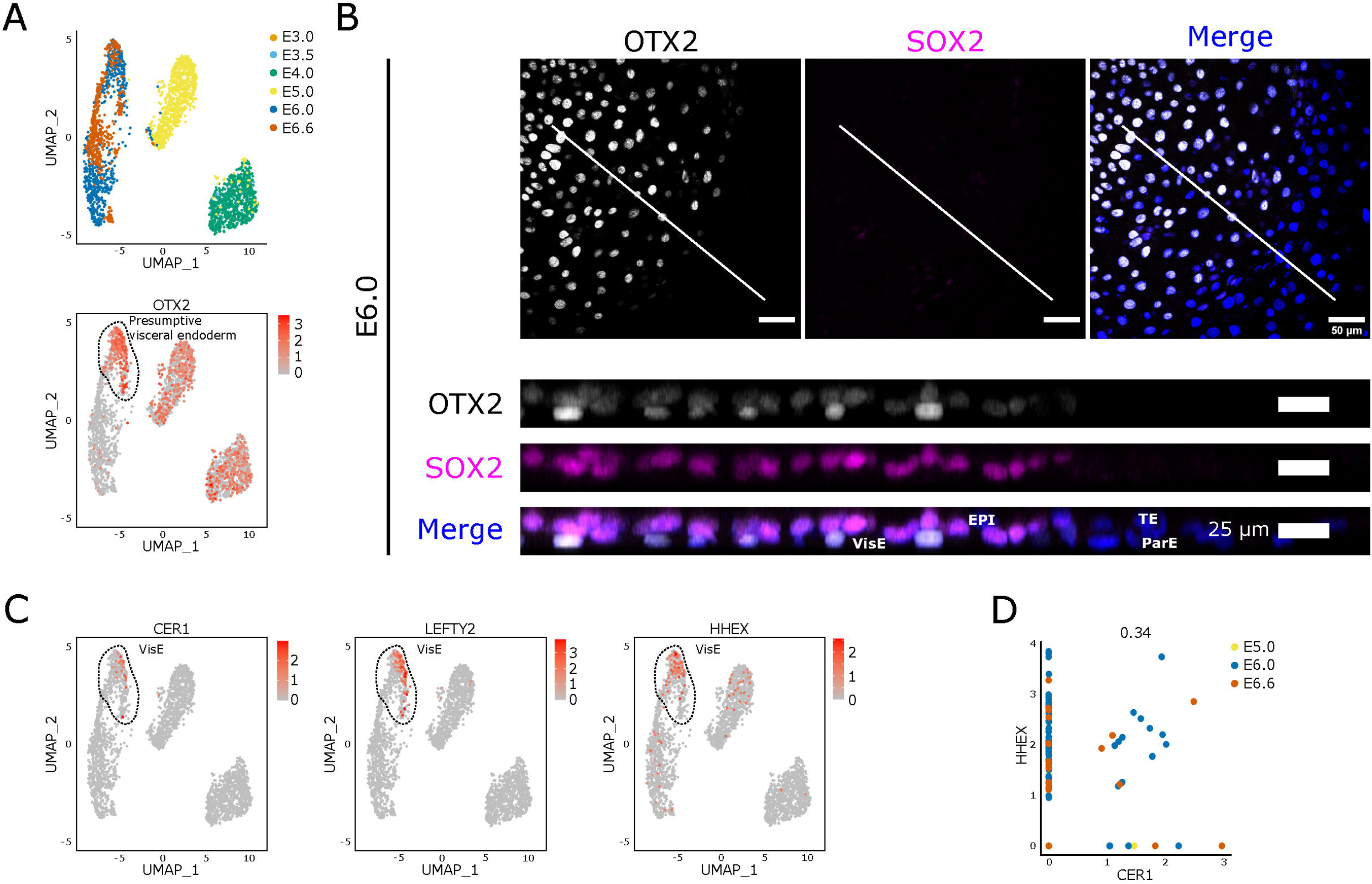
Polarisation of the visceral endoderm. (**A**) Two-dimensional UMAP representation of 2,777 single-cell transcriptomes with only PE cells, showing embryonic days (upper panel) or *OTX2* expression (lower panel). (**B**) Immunofluorescence labelling of epiblasts (E6.0; n = 7) with OTX2 and SOX2 antibodies. Merge pictures include DAPI staining. The white lines in the upper panels (scale bars: 50 μm) indicate the position of the optical sections shown in the lower panels (scale bars: 25 μm). (**C**) Two-dimensional UMAP representation of the 2,777 PE single-cell transcriptomes with expression of anterior VisE markers *CER1, LEFTY2* and *HHEX*. (**D**) Scatter plot representation of single cells expressing *CER1* and *HHEX* in the ParE and VisE clusters. Colours indicate embryonic days.

To determine the timing of anterior visceral endoderm formation, we next examined the expression of *CER1, HHEX* and *DKK1* (**Fig. 3C**), and identified a sub-cluster of VisE cells co-expressing *HHEX* and *CER1* at E6.0 (**Fig. 3D**). These findings point to the onset of VisE polarization at E6.0 in rabbit. Remarkably, we noticed the upregulation of *BMP2* and *BMP4* in the parietal endoderm (**Fig. S3A**) as previously described using RNA-FISH (Hopf et al. 2011).

To characterize mesodermal *vs*. endodermal segregation, we used our rabbit/mouse/cynomolgus integrated dataset (**Fig. S2**) and identified anterior (EPI_ant) and posterior (EPI_post) epiblast within the EPI cells (**Fig. 4A**). We also identified a minor cluster located between these EPI_ant and Epi_post clusters that we qualified as EPI_int (EPI_intermediate). We then performed a principal component analysis (PCA) on the EPI_post cells (n = 239) using 15 markers (5 per tissue: anterior epiblast, endoderm and mesoderm; **Fig. 4B**). Cells formed two separate clusters, with one cluster having higher expression of mesodermal markers, including *TBXT, HAND1, LEF1, PDGFRA*, and *WNT5A*, and the other cluster having higher expression of definitive endoderm markers, including *CHRD*, *CER1*, *GSC*, *OTX2*, and *HHEX* (**Fig. 4B**). To localize primordial germ cells (PGCs) in the embryos, we performed immunofluorescence analysis to identify cells positive for both TFAP2C and TBXT (**Fig. S3B**). These double positive cells (arrows in **Fig. S3B**) were observed only in the posterior epiblast at E6.6, not at E6.0. These observations are in line with previous studies reporting segregation between mesoderm and endoderm after E6.0 in rabbit embryos, followed by the emergence of PGCs at E6.6 (Hassoun et al., 2009a; Hassoun et al., 2009b; Hopf et al., 2011; Kobayashi et al., 2021; Viebahn et al., 2002).

**Figure 4:**
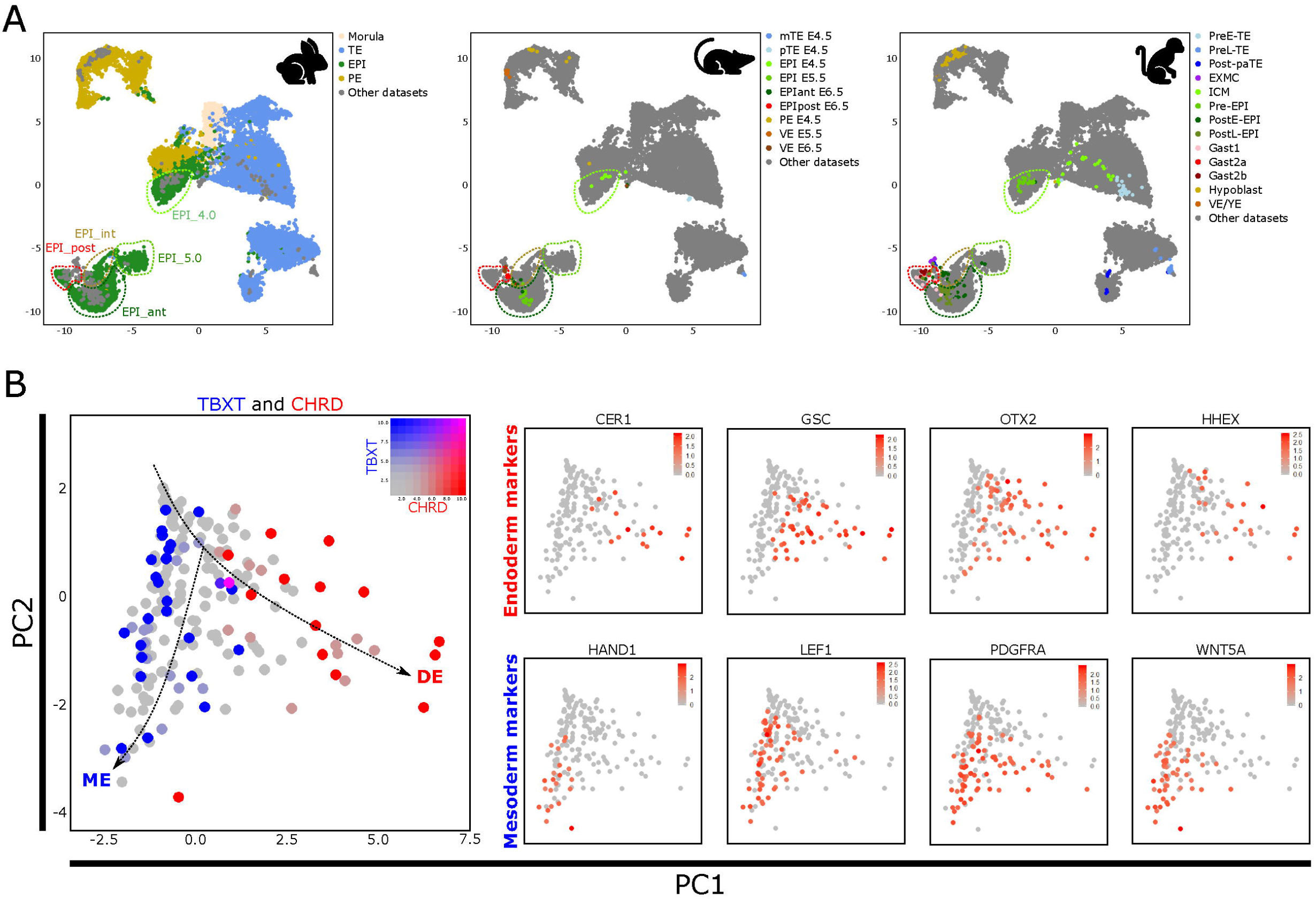
Identification of late epiblast subclusters and onset of gastrulation. (**A**) UMAP representation of the integrated transcriptome dataset generated by merging the rabbit, mouse and cynomolgus monkey datasets as shown in Fig. S2. The dashed lines highlight samples from posterior/gastrulating epiblast cells in red, anterior epiblast cells in dark green, and early epiblast cells in light green. (**B**) Two dimensional PCA of the transcriptome of the 237 EPI_post cells showing mutually exclusive markers of definitive endoderm (DE; *CHRD*, *CER1*, *GSC*, *OTX2* and *HHEX*) and mesoderm (ME; *TBXT*, *HAND1*, *LEF1*, *PDGFRA*, *WNT5A*).

### Gradual alteration of the transcriptome in the pluripotency continuum

To characterize the transcriptomic changes taking place in the pluripotency continuum of rabbit embryos, we focused subsequent analysis on morula, ICM, and EPI cells within the 10x dataset. The resulting “PLURI dataset” included 4,315 cells from six developmental stages. We then examined the expression of known markers of naive, formative and primed pluripotency (**Fig. 5A, Table S2)**. A stronger expression of naive pluripotency genes was observed in morula, ICM, and EPI_4.0/5.0 cells, whereas a stronger expression of formative genes was observed in EPI_5.0, EPI_int and EPI_ant cells. Finally, primed pluripotency-associated genes were mostly up-regulated in the EPI_post cells. These results suggest that the epiblast cells of rabbit embryos between E3.5 and E6.6 encompasses the whole pluripotency continuum.

**Figure 5:**
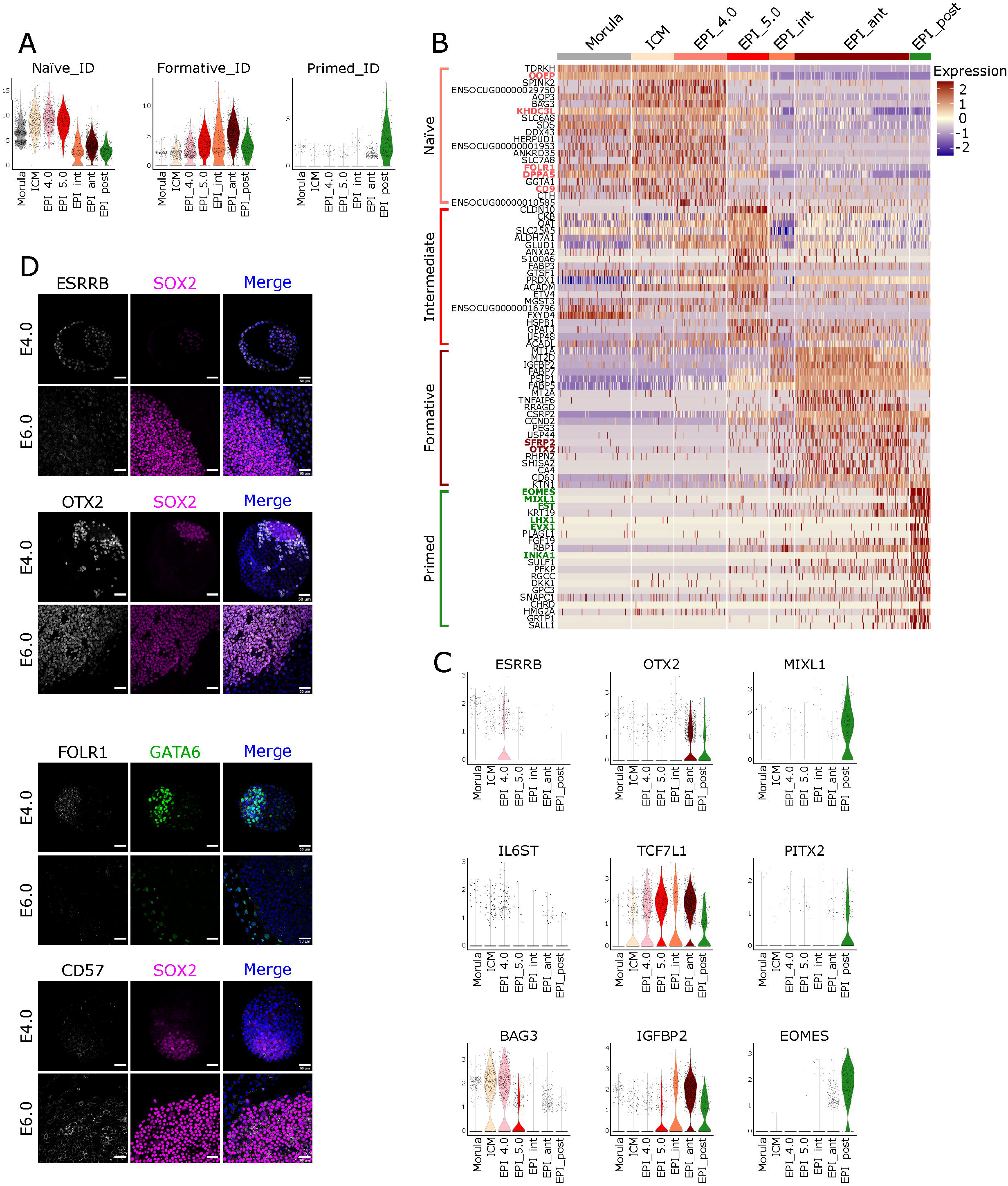
Transcriptome map of the pluripotency continuum. (**A**) Violin plots representing the sum expression of genes associated with the naive (*CDH1, IL6ST, OOEP, PRDM14, RIF1, TFAP2C*, *TFCP2L1*, *ZFP42*, *TBX3* and *ESRRB*), formative (*OTX2*, *TCF7L1*, *ETV5*, *SALL2*, *SOX3*, *SOX11*, *DNMT3A*, *DNMT3B*, *CRABP2* and *POU3F1*) and primed (*MIXL1*, *PITX2*, *TBXT, TET3, FGF10, WNT5A, FOXA2, CDH2, CER1* and *GATA4*) state of pluripotency. (**B**) Heatmap representation grouping 20 markers (from **A**) with the 20 most differentially expressed genes (10x Genomics “PLURI dataset”) in EPI_4.0 *vs*. all, EPI_5.0 *vs*. all, EPI_ant *vs*. all, and EPI_post *vs*. all. the other cell types. (**C**) Violin plot representation of gene expression calculated from the “PLURI dataset.” 1^rst^, 2^nd^ and 3^rd^ rows show expression of naive, formative and primed pluripotency gene markers, respectively. (**D**) Immunofluorescence detection of ESRRB, SOX2, OTX2, FOLR1, GATA6, and CD57 in E4.0 (n = 23) and E6.6 (n = 16) embryos. Merge pictures include DAPI staining. Scale bars: 50 μm.

A heatmap was generated with the 20 most overexpressed genes in each cell type. Morula, ICM and EPI_4.0 cells had a higher expression of naive pluripotency markers including *ESSRB, DPPA5, OOEP, BAG3* and *FOLR1* (**Fig. 5B,C, Table S2**). These markers declined in EPI_5.0, concomitantly with the upregulation of a new set of markers, including genes associated with the exit of naive pluripotency *ETV4* and *TCF7L1*. Other markers enriched in EPI_5.0 included cell-cell interactions- and metabolic pathways-related genes including *CLND10, CKB, OAT, ALDH7A1, ANXA2*, and *S100A6* (**Fig. 5B,C)**. A set of genes including formative/primed pluripotency marker *OTX2* was up-regulated in both EPI_ant and EPI_post cells (**Fig. 5B,C**). Finally, several genes were up-regulated only in the EPI_post cells, including *MIXL1, PITX2* and *EOMES*, revealing the onset of gastrulation. Differential localization of DPPA5, ESRRB, OTX2 and FOLR1 was confirmed by immunofluorescence (**Figs 5D, S4A**); ESRRB and FOLR1 were more strongly detected in the EPI at E4.0 compared to E6.0 embryos, whereas OTX2 was more strongly detected in the EPI at 6.0 than in the EPI at 4.0. The primed pluripotency marker CD57 was also strongly detected in the EPI at 6.0. Genes up-regulated in epiblast cells (EPI_ant and EPI_post) also included metabolic pathways-related genes (i.e. *MT1A*, *MT2D, FABP5*, and *FABP7*; **Fig. 5B**). Together, these results suggest alterations in metabolic pathways concomitant with the transition from the naive to primed pluripotency state. This observation was corroborated by GO and KEGG pathway enrichment analysis based on the 500 most differentially expressed genes between early and late epiblast cells (**Fig. S4B,C, Table S2**). Taken together, these results allowed us to identify gene markers of the naive, formative and primed pluripotency states in rabbit preimplantation embryos.

### Modification of the epigenetic landscape in the pluripotency continuum

We investigated the DNA methylation dynamics during rabbit preimplantation development using immunodetection of 5-methylcytosine (5meC) and 5-hydroxy-methylcytosine (5hmeC) in E3.0 to E6.6 embryos (**Fig. 6A**). 5meC fluorescence was low in morulae (E3.0) and early blastocysts (E3.5) compared to later stages. There was also significantly higher 5meC fluorescence in EPI cells compared to TE cells in E6.0 and E6.6 embryos. On the other hand, 5hmC signal was higher in early stages (E3.0/E3.5), suggesting active demethylation. At later stages, the signal progressively disappeared in the TE cells and then in the epiblast. Finally, it could again be detected at E6.0 and E6.6 in EPI cells. These 5meC and 5hmeC fluorescence patterns are consistent with the increasing expression of DNA methyltransferases (DNMTs) and the decreasing expression of ten-eleven translocation methylcytosine dioxygenases *TET1* and *TET2*, observed during the transition from ICM to EPI (**Fig. 6B**). Note, however, that *TET3* expression increased between the ICM and late EPI stages, which may explain why 5hmeC is still detected in the EPI at E6.0 and E6.6, despite the increased level of 5meC and increased expression of *DNMTs*. Thus, the rabbit embryo shows DNA methylation dynamics similar to those previously described in mice and humans (Zhang et al., 2018; Zhu et al., 2018).

**Figure 6:**
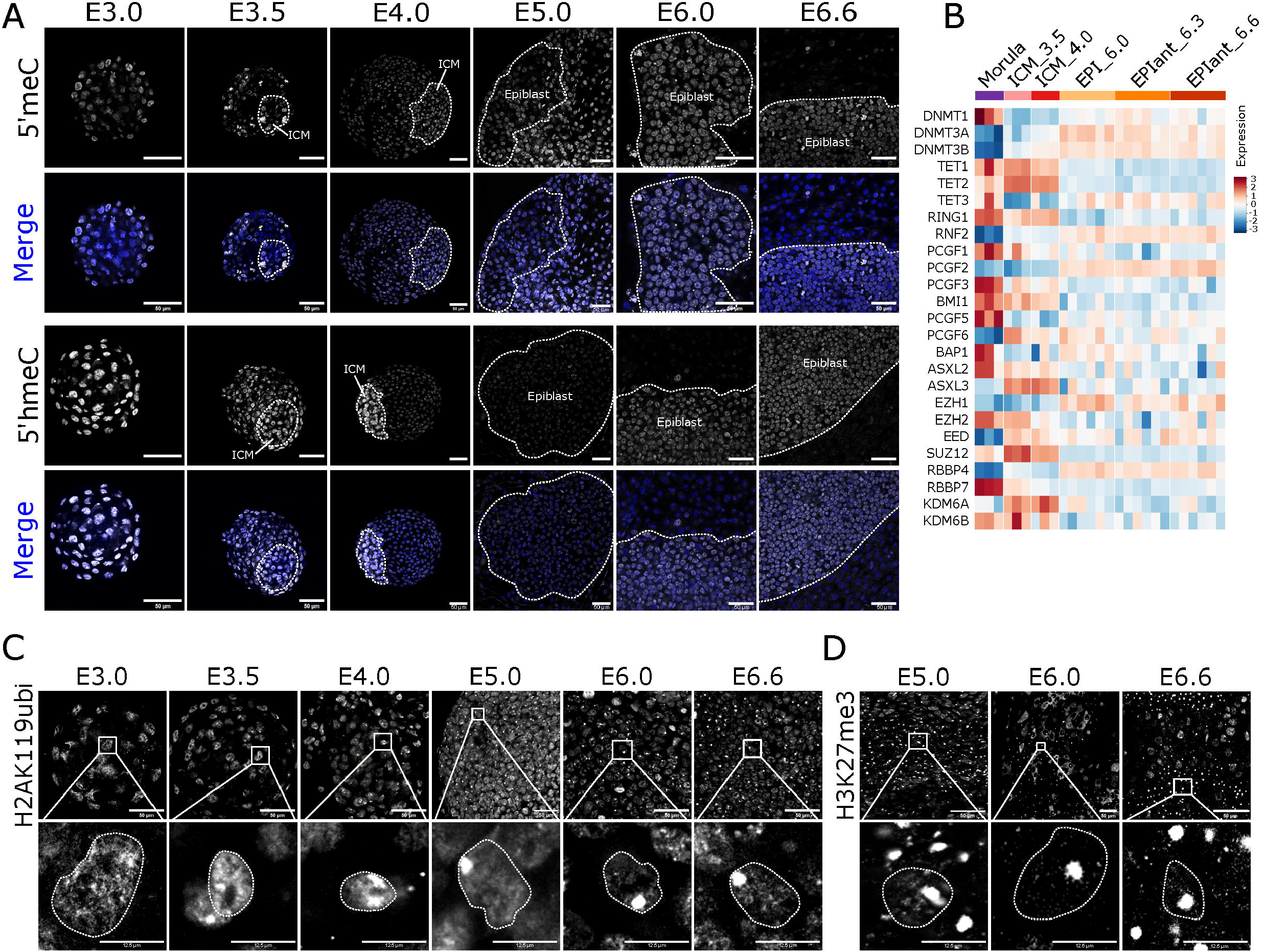
Epigenetic characterization of the pluripotency continuum. (**A**) Immunofluorescence detection of 5-methylcytosine (5meC) and 5-hydroxymethylcytosine (5hmeC) in E3.0 morula (n = 16), E3.5 (n = 18), E4.0 (n = 22) and E5.0 (n = 25) blastocysts, E6.0 (n = 6) and E6.6 (n = 5) epiblasts. ICM and epiblast are surrounded by dotted lines. Merge images include propidium iodide staining in blue. Scale bars: 50 μm. (**B**) Heatmap representation of bulk RNAseq dataset showing the expression of 25 genes involved in DNA methylation and polycomb repressive complexes. (**C**) Immunolabelling of histone 2A lysine119 ubiquitinated (H2AK119ubi) marks in embryos at E3.0 to E6.6 (n = 41) (scale bar: 50μM). Enlargements show single nuclei (scale bar: 12.5 μm). (**D**) Immunolabelling of histone 3 lysine 27 tri-methylation (H3K27me3) marks in embryos at E5.0-E6.6 (n = 12) (scale bar: 50 μm). Enlargements show single nuclei (scale bar: 12.5 μm).

Through the study of X chromosome coating by *XIST* RNA, it was shown that X inactivation begins at the morula stage in the rabbit (Okamoto et al., 2011). To describe this process in more detail across the pluripotency continuum, we analysed H2AK119ub and H3K27me3 marks, two post-translational modifications of histones associated with X chromosome inactivation in mice (Chaumeil et al., 2011). At the morula stage, H2AK119ub immunostaining appears as small diffuse nuclear spots (E3.0) (**Fig. 6C**). From the early blastocyst stage onward, labelling begins to form foci in half of the embryos analysed (n = 34). In those, the percentage of cells with a single nuclear focus increased from 15% (E3.5) to 100% (E5.0). Immunostaining of the repressive mark H3K27me3 revealed a similar dynamic (n = 12 per stage) (**Figs 6C, S5A**). Consistent with these observations, the bulk RNAseq dataset revealed that expression of the H2AK119ub erasers, *ASXL2* and *ASXL3*, as well as expression of the H3K27me3 erasers, *KDM6A* and *KDM6B*, were down-regulated during the transition from ICM to EPI_6.0/6.3/6.6 (**Fig. 6B**). From these results, we conclude that X chromosome inactivation by repressive marks begins as early as E3.5 and is established at E5.0.

### Gradual switch from OXPHOS to glycolysis-dependent metabolism in the pluripotency continuum

The bulk RNAseq dataset was used to investigate the expression of genes associated with oxidative phosphorylation (OXPHOS) and glycolysis. A stronger expression of both nuclear (NC)- and mitochondrial (MT)-encoded genes related to OXPHOS metabolism was observed in TE and ICM cells compared to EPI_6.0/6.3/6.6 (**Fig. 7A**). Analysis of the single-cell “PLURI dataset” confirmed this observation, showing a gradual downregulation of *NDUFV2, UQCRQ, COX8A*, and *ATP5PD* gene expression between E3.0 and E6.6 (**Fig. 7B**). In contrast, *LDHA, LDHB*, and *PKM*, which are key genes of glycolysis, were expressed at much higher levels in EPI_6.0/6.3/6.6 compared to ICM (**Fig. 7A**). The single-cell “PLURI dataset” confirmed this finding, showing a gradual increase in the expression of these genes between E3.0 and E6.0 (**Fig. 7B**). These observations are consistent with a switch from OXPHOS to glycolysis for energy production between early blastocysts and pre-gastrula stage embryos in rabbits, which correlates with observations made in mice (Houghton, 2006). However, other genes involved in glucose uptake and metabolism, including *SLC, HK*, and *PDH*, were already expressed at early embryo stages (ICM and E3.0 to E4.0), which suggests that the pluripotent cells of the ICM and early EPI are poised for upregulating glucose metabolism-based energy production.

**Figure 7:**
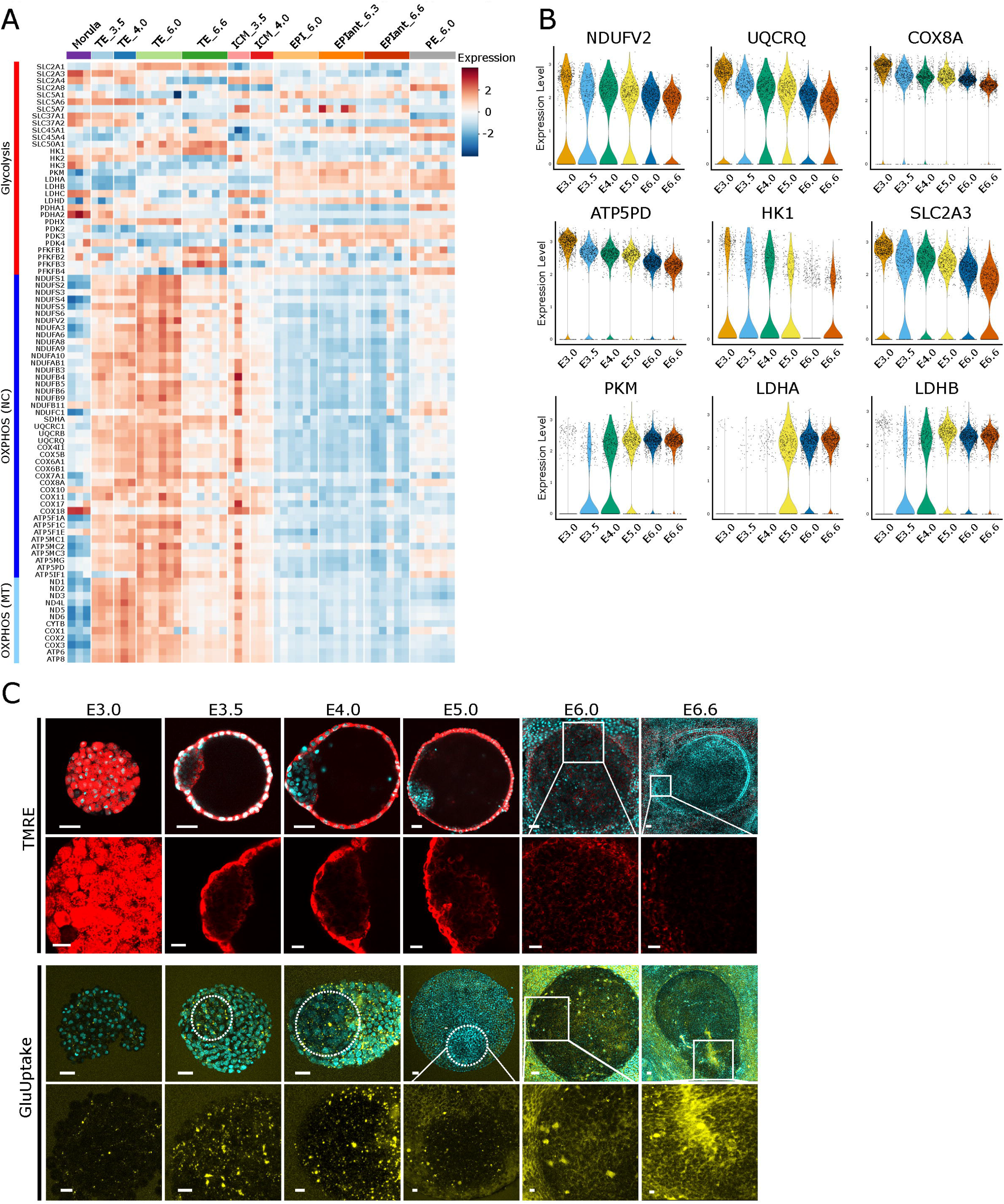
Metabolic characterization of the pluripotency continuum. (**A**) Heatmap representation of the bulk RNAseq dataset and showing the expression of genes involved in glycolysis and oxidative phosphorylation (OXPHOS). NC, nuclear genome; MT mitochondrial genome in morula (E2.7), TE (E3.5-E6.6), ICM (E3.5/E4.0), EPI (E6.0-E6.6) and PE_6.0. (**B**) Violin plot representation of the expression of genes involved in OXPHOS and glycolysis at different embryonic stages in the “PLURI dataset”. (**C**) Metabolism analysis of live rabbit embryos between E3.0 and E6.6. The 1^st^ vertical row show images of embryos (n = 34, 5 to 11 embryos per stage) treated with both tetramethylrhodamine ethyl ester (TMRE, red staining) and Hoechst (blue). The 2^nd^ vertical row shows enlargements of ICM/epiblast cells after TMRE staining. The 3^rd^ vertical row show images of embryos (n = 90, 5 to 19 embryos per stage) treated with GluUptake (yellow staining) and Hoechst (blue). The 4^th^ vertical row shows enlargements of ICM/epiblast cells after GluUptake staining. Scale bars: 50 μm, and 20μm for enlargements.

To investigate mitochondrial activity during rabbit preimplantation embryo development, embryos between E3.0 and E6.6 were treated with tetramethylrhodamine ethyl ester (TMRE), which reveals mitochondrial membrane depolarization (ΔΨm). Morula and TE cells showed strong TMRE labelling, indicating high ΔΨm and strong OXPHOS activity (**Fig. 7C**). TMRE labelling was much lower in the ICM and EPI cells of E3.5 to E6.0 embryos, and became undetectable by E6.6. This result was confirmed by CellROX assay, which labels the reactive oxygen species (ROS) produced by OXPHOS (**Fig. S5B**). These results are consistent with the expression pattern of *NDUF*, *COX*, *UQCR*, and *ATP5* families of genes, which are highly expressed in TE cells, and have low expression in EPI_6.0/6.3/6.6 cells. Fluorometric determination of 2-deoxyglucose incorporation was performed to study glycolysis in rabbit embryos (**Fig. 7C**). Weak 2-deoxyglucose fluorescence was observed in morulae (E3.0), but gradually increased in later stages (from E3.5 onwards), first in TE cells and then in EPI cells. Strong fluorescence was observed in the primitive streak region in E6.6 embryos, correlating with the higher expression of *LDHA* and *PKM* observed in transcriptome studies (**Fig. 7B**).

### Robust markers of naive pluripotency in rabbits are common to either mice or primates and rarely to both

We sought to further characterize the naive pluripotency state in rabbit embryos and compare to current data in mice and primates. To address this, we analysed the differentially expressed genes between ICM and TE, ICM and PE_6.0, and ICM and EPI_6.0/6.3/6.6 in the bulk RNAseq datase, which resulted in 1,260 up-regulated genes in ICM cells compared to the other lineages (p < 0.01) (**Figs 8A, S6A, Table S3**). We performed a similar differential analysis between ICM cells (E3.5/E4.0) compared to other cell types (E3.5-E6.6) in the 10x-genomics dataset (average log Fold change > 0.25), and identified 161 up-regulated genes in ICM cells compared to the other lineages (**Fig. 8B, Table S4**). We found 95 genes in common between those two sets of up-regulated genes (**Fig. 8C, Table S4**). Among these 95 genes were pluripotency-associated genes *BAG3, DPPA5, SOX15*, chromatin regulator-encoding genes *KDM4A, KMT2C, SMARCAD1*, and metabolic pathways associated genes such as *ALDH7A1* (**Fig. 8D, Table S4**). This observation was corroborated by GO analysis that revealed enrichment for DNA metabolic processes and “DNA repair” (**Fig. 8E, Table S4**). Notably, both a higher level of *KDM4A* transcripts at E4.0 and ICM-specific detection of KDM4A protein were observed (**Fig. 8F,G**). We thus identified key markers of naive pluripotency that might be useful to characterize embryo-derived and induced pluripotent stem cell (ESC and iPSC) lines in the rabbit.

**Figure 8:**
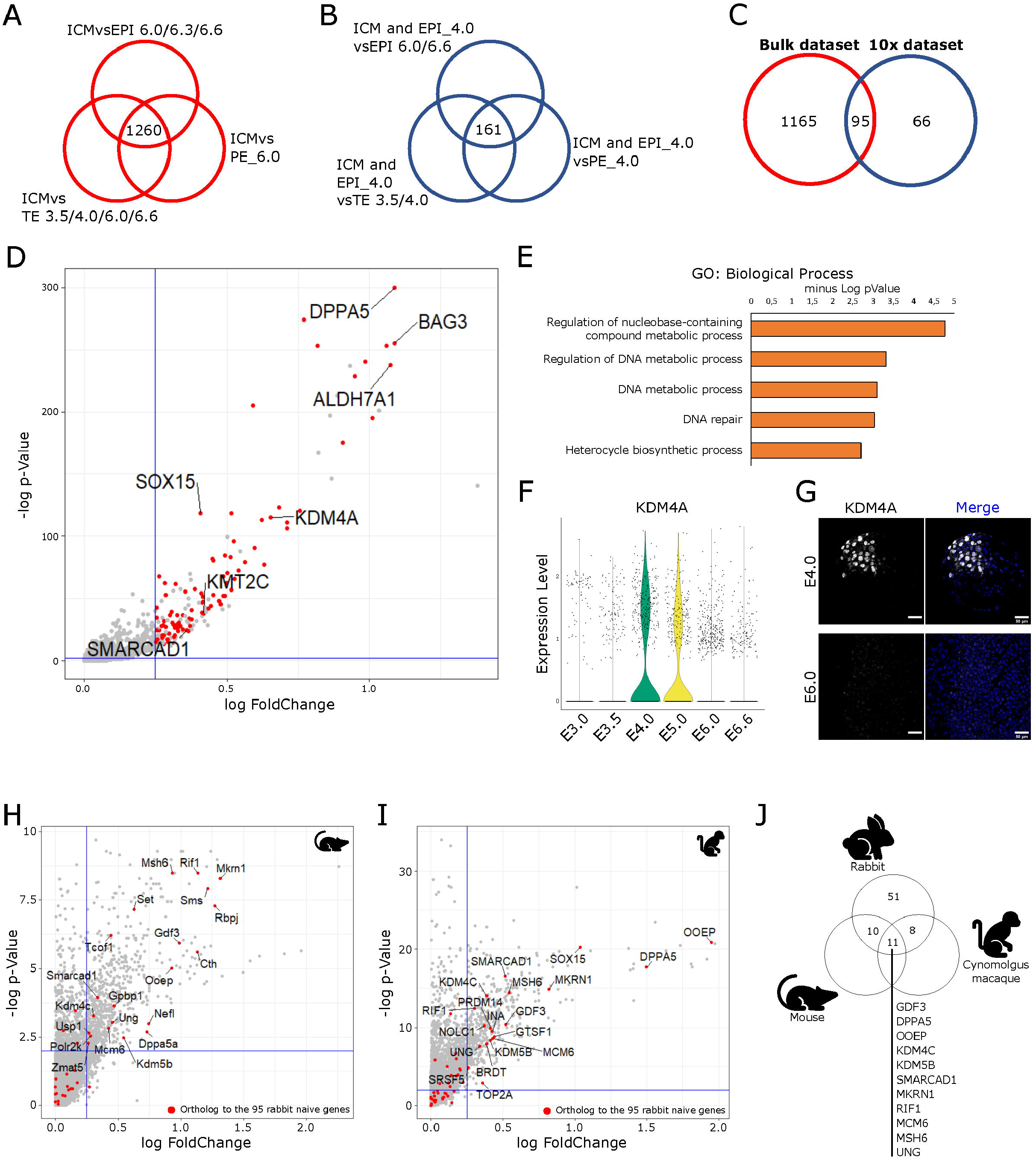
Defining markers of naive pluripotency in rabbits. (**A**) Venn diagram representing the total number of up-regulated genes between ICM (E3.5/E4.0) and TE (E3.0-E6.6), EPI (E6.0-E6.6) or PE_6.0 in the bulk RNAseq dataset. (**B**) Venn diagram representing the total number of up-regulated genes between ICM/EPI_early (E3.5/E4.0) and TE_3.5, TE_4.0, TE_int1, TE_int2 (E3.0-E4.0); EPI_ant, EPI_int, EPI_post (E6.0-E6.6); or PE_4.0 (E4.0) in the 10x RNAseq dataset. (**C**) Venn diagram representing the intersection between the genes up-regulated in the ICM (E3.5/E4.0) and ICM/ EPI_4.0 (E3.5/E4.0) cells according to the bulk (A) and 10x (B) RNAseq data. (**D**) Volcano plot representations of the differentially expressed genes between the ICM/EPI_4.0 (E3.5/E4.0) and TE_3.5, TE_4.0, TE_int1, TE_int2 (E3.0-E4.0); EPI_ant, EPI_int, EPI_post (E6.0-E6.6); or PE_4.0 (E4.0) in the 10x RNAseq dataset (see fig. 8B). In red are the 95 genes commonly enriched in the naive cells of the 10x and Bulk datasets (see fig. 8C) (**E**) Gene Ontology biological process analysis of the 95 up-regulated genes in the ICM/EPI cells of rabbit embryos, identified in **C**. (**F**) Violin plot representation of *KDM4A* expression in morula/ICM/EPI cells at E3.0 to E6.6 (“PLURI dataset”). (**G**) Immunofluorescence detection of KDM4A in E4.0 (n = 4) and E6.0 (n = 4) rabbit embryos and counterstained with DAPI. Scale bars: 50 μm. (**H**) Dot plot representation (-log p-value as a function of log fold-change) of the positively differentially expressed genes between ICM (E3.5/E4.0) and other cell types in mice (Argelaguet et al., 2019). The 81 ortholog genes are shown in red. Among them, only those that are significantly enriched in naive mouse cells are named. (**I**) Dot plot representation (-log p-value as a function of log fold-change) of the positively differentially expressed genes between ICM/EPI (E6.0/E9.0) and other cell types in cynomolgus monkey (Nakamura et al., 2016). The 81 ortholog genes are shown in red. Among them, only those that are significantly enriched in primate cells are named. (**J**) Venn diagram representing the distribution of the 95 ICM/EPI-up-regulated genes identified in (C), compared to mice and cynomolgus monkeys naive markers.

We then further examined the JAK-STAT, NOTCH, WNT, MAPK, and TGF-beta signalling pathways, all of which are associated with the regulation and maintenance of pluripotency in rodents and primates (Bayerl et al., 2021; Boroviak et al., 2015; Boroviak et al., 2018; Mohammed et al., 2017; Nakamura et al., 2016). Consistent with naive pluripotency data in mice (Mohammed et al., 2017), the expression of components of the JAK-STAT signalling pathway, including *IL6R*, *IL6ST*, *IL4R*, *STAT4* and *STAT3*, were increased in ICM cells versus TE, PE and late EPI cells (**Fig. S6B**). Notably, the expression of *NOTCH* and NOTCH target gene *HES1* were low, whereas the expression of the NOTCH ligand *JAG1* and the NOTCH inhibitor *NUMB* were high in ICM cells compared with other cells (**Fig. S6C**). Low NOTCH activity in rabbit ICM is consistent with recent data indicating that treatment of human PSCs with the NOTCH pathway inhibitors, DBZ and RIN1, enhances naive pluripotency (Bayerl et al., 2021). Overall, these results suggest that the mechanisms controlling naive pluripotency in rabbit preimplantation embryos utilize signalling pathways that function in mice, primates, or in both.

Finally, we asked which of the aforementioned 95 genes are also specific to the ICM and early EPI cells in both mice and cynomolgus macaques, making use of the single-cell RNAseq data previously generated (Nakamura et al., 2016). Since we could not find any orthologs for 14 of the 95 genes, we had to narrow down the analysis to 81 genes (**Table S4**). In the mouse and cynomolgus macaque datasets, ICM and early EPI cells were identified using landmark markers of naive pluripotency in mice (*Fgf4, Esrrb* and *Klf2*) (**Fig. S7A**) and cynomolgus macaques (*FGF4, KLF17*, and *SOX15*) (**Fig. S7B**). The most overexpressed genes between ICM and early EPI cells compared to all other cell types were identified and represented by heatmaps (**Fig. S7A,B, Table S4**). Genes more highly expressed in naive cells were represented by volcano plots, highlighting the 81 ortholog genes (**Fig. 8H,I**). Of these 81 genes, 10 were also up-regulated in mice (average log fold-change > 0.25) and 8 were up-regulated in cynomolgus macaques (average log fold-change > 0.25). These genes are associated with “pluripotency regulation,” “chromatin organization”, “mRNA splicing” and “DNA metabolism and repair” as shown in an interaction map of biological processes (**Fig. S7C**). Finally, a comparison of the naive markers in the three species led to the identification of 11 common genes: *GDF3, KDM4C, KDM5B, SMARCAD1, RIF1, MSH6, MCM6, UNG, DPPA5, OOEP* and *MKRN1* (**Fig. 8J**). Taken together, these results show that rabbit preimplantation embryos share some of the marker genes for naive pluripotency with those of mice, and some with those of cynomolgus macaques.

## Discussion

Our study describes the results of the first thorough investigation of the rabbit preimplantation embryos at the single-cell level. It combines bulk and single-cell RNA sequencing, protein immunolabelling, and fluorometric quantification to characterize the transcriptome, epigenome and metabolome of the pluripotent cells from the morula to early gastrula stage. Four main findings emerge from this study: (1) The three early lineages, EPI, PE and TE are fully segregated between E5.0 and E6.0. (2) ICM and early EPI cells (E3.5/E4.0) and late EPI cells (E6.0/E6.6) exhibit the cardinal features associated with naive and primed pluripotency, respectively; E5.0 is a transitional stage in that respect. (3) Novel markers of naive pluripotency were identified, including *MKRN1* and *OOEP*; (4) Although the rabbit is evolutionarily closer to mouse than to primates, the transcriptome of rabbit pluripotent cells shares many common features with that of humans and non-human primates, including markers of naive pluripotency.

In Eutherian mammals, TE formation is triggered by the asymmetric segregation of keratins and polarization of the outer cells of the morula at the onset of compaction (Lim et al., 2020 Nature; Gerri et al., 2021). In our study, the early expression of *KRT8, KRT18, GATA2*, and *GATA3* indicates an early onset of the TE program in some morula cells. In contrast, *FABP3* expression increases later, at E4.0, in the TE of the expanded blastocyst. *OCT4*, initially expressed in all cells of the early blastocyst, including the TE, becomes restricted to EPI cells only at E5.0, consistent with a previous study (Canon et al., 2018). These results point to the progressive differentiation of TE lineage. It is also from E4.0 onwards that the expression of *SOX2* and *GATA6*, markers of EPI and PE, respectively, become mutually exclusive and the two cell types separate to form two distinct cell compartments at E5.0, consistent with a previous study (Piliszek et al., 2017). These results indicate that the late segregation of EPI, TE, and PE lineages is similar to what has been described in human and non-human primates (Meistermann et al., 2021; Nakamura et al., 2016; Stirparo et al., 2018).

Studying the transition from the morula stage to gastrulation is particularly challenging in many species as the developmental stage at which it occurs corresponds roughly to the time when embryos implant in the uterus. This is particularly true of primates, where access to the newly implanted embryo (between E8 and E13) is virtually impossible. In contrast, rabbit embryos are well suited for investigating this transition because they have delayed embryo implantation until after the onset of gastrulation (Fischer et al., 2012; Puschel et al., 2010). We found that the epiblast of E5.0 rabbit embryos is a transition state, characterized by the downregulation of naive pluripotency markers and the upregulation of formative and primed pluripotency markers. In addition, it is at this stage that we observed the shift to predominantly glycolic metabolism, genome re-methylation, and X chromosome inactivation, all of which are considered as key events of the naive-to-primed state transition in rodents and primates (Davidson et al., 2015; Devika et al., 2019). Despite embryo staging prior to cell preparation, some overlap between naive, formative and primed markers can be observed, which may reflect asynchrony of gene expression dynamics. This heterogeneity emphasizes the transitional-stage notion associated with E5.0 rabbit embryos.

Our study identified new marker genes for naive pluripotency in rabbit, a species in which it has not yet been possible to derive truly naive pluripotent stem cell lines (Osteil et al., 2016; Tapponnier et al., 2017). Eleven of these genes are also specifically expressed in the ICM and early EPI in mouse and cynomolgus macaque embryos. *GDF3* and *SMARCAD1* are well-known pluripotency-associated genes. *GDF3* encodes a ligand of TGFβ superfamily that blocks BMP signalling and regulates cell fate in stem cells and embryos (Levine and Brivanlou, 2006). *SMARCAD1* encodes a SWI/SNF-like chromatin remodeller with ATPase activity. Its inactivation is detrimental to mouse ESC pluripotency (Sachs et al., 2019; Xiao et al., 2017). Three other genes have also functional characteristics expected of a naive pluripotency regulator. *MKRN1* is a target of OCT4 (Cassar et al., 2015), and encodes an E3 Ubiquitin Transferase involved in the degradation of p53 and p21 and promotes cell-cycle progression (Lee et al., 2009). *DPPA5* encodes an RNA binding protein that interacts with mRNAs encoding pluripotency and cell-cycle regulators (Tanaka et al., 2006). Moreover, DPPA5 binds NANOG and enhances its function while also preventing its degradation (Qian et al., 2016). *OOEP* belongs to the same family as *DPPA5* (Pierre et al., 2007) and known to be expressed in the ICM cells and naive PSCs in various species (Messmer et al., 2019; Mohammed et al., 2017; Nakamura et al., 2016; Stirparo et al., 2018). *MKRN1, DPPA5* and *OOEP* are therefore new markers of naive pluripotency. Our study also highlighted the potential role of histone lysine demethylases in naive pluripotency. Three of them are specifically expressed in the ICM and early EPI of rabbit embryos: *KDM4A, KDM4C* and *KDM5B*. These enzymes “erase” methylation on histone H3K9 and H3K4 respectively. They have been shown to co-occupy promoters in mouse ESCs (Tripathi et al., 2021), suggesting a functional link to gene expression programmes. Finally, upregulation of *MCM6, MSH6, RIF1* and *UNG* also characterizes naive pluripotency based on our inter-species comparison. These four genes are related to DNA repair and replication, two pathways known to be involved in maintaining the genome integrity and identity of pluripotency cells (Mason et al., 2009).

Although many of the naive and primed pluripotency markers identified in rabbits are common to mice and primates, our study also highlights some noticeable differences. The IL6 receptor-encoding gene *IL6R* is expressed in the ICM of rabbit and primate embryos, but is replaced by the LIF receptor-encoding gene *LIFR* in mice (Boroviak et al., 2018). *SOX15* is a naive pluripotency marker in rabbits (this study) and primates, but not in mice (Stirparo et al., 2018); (Boroviak et al., 2018). *TDGF1* is expressed in ICM and early EPI cells but down-regulated in late EPI cells in mice (Boroviak et al., 2015; Mohammed et al., 2017), whereas it is only expressed in late EPI in rabbit and cynomolgus macaque (Nakamura et al., 2016). Thus, these genes exhibit an opposite expression pattern in rabbits (this study) and cynomolgus macaques *vs*. mice. These interspecies comparisons clearly suggest a similarity between rabbits and primates in the expression of pluripotency regulators across the pluripotency continuum. Arguably, this makes the rabbit a suitable species for studying the embryo colonization capacity of human naive PSCs and the generation of inter-species chimeras (Aksoy et al., 2021).

## Materials and Methods

### Production and dissection of rabbit embryos

All procedures in rabbits were approved by the French ethics committee CELYNE (approval number APAFIS#6438 and APAFIS #2180-2015112615371038v2), and COMETHEA n°45, registered under n°12/107 and n°15/59. Sexually mature New Zealand white rabbits were injected with follicle-stimulating hormone and gonadotropin-releasing hormone, followed by artificial insemination as previously described (Teixeira et al., 2018). Embryos were flushed from explanted oviducts 65–159 h after insemination.

For bulk RNAseq, morulae were collected at embryonic day 2.7 (E2.7), pooled by ten and immediately dry-frozen. Blastocysts were collected at E3.5 and E4.0. Mucin coat and zona pellucida (ZP) were removed by protease digestion (Sigma P8811-100MG). Zona-free embryos were subsequently cultured in TCM199 supplemented with 10% foetal bovine serum (FBS, Sigma M4530) until they regain a normal morphological aspect. Inner Cell Masses (ICMs) were separated from the trophectoderm (TE) by moderate immune-surgery: briefly, blastocysts were incubated in anti-rabbit whole goat serum (Sigma R-5131) at 37°C for 90 min, washed thoroughly and then incubated with guinea pig complement serum (Sigma S-1639) for 5 min. After washing in PBS supplemented with 10% FBS, ICMs were mechanically dissociated from the TE by gentle pipetting with a glass pipette. Samples (ICM and TE) were pooled by ten and immediately dry-frozen. At E6.0, the mucoprotein layer was removed mechanically using glass microcapillaries. Zona-free embryos were opened and flattened on a plastic dish in FHM medium to expose the embryoblast with the primitive endoderm on top. The primitive endoderm (visceral part) was first dissociated by careful scratching with a glass needle, and the epiblast was then separated from the trophectoderm with a microscalpel. Samples were dry-frozen. At E6.3 and E6.6, embryos were processed as described for E6.0 embryos. The anterior part of the epiblast was isolated from the posterior epiblast by manual micro-dissection prior to dry-freezing. For E6.3 embryos, the procedure for isolating the anterior epiblast was validated *a posteriori* by analysing *TBXT* expression by RT-qPCR.

For 10x single-cell RNAseq, embryos collected at E3.0, E3.5, E4.0 and E5.0 were treated with protease from Streptomyces griseus (Sigma-Aldrich P8811) at 37°C followed by mechanical dissociation with glass microcapillaries to remove the mucin coat and ZP. For embryos collected at E6.0 and E6.6, mucoprotein layers were removed mechanically with forceps. The embryonic disks were then dissected mechanically with forceps and pooled prior to single-cell dissociation. For cell singularization, E3.0, E3.5, E4.0 and E5.0 embryos were treated with 0.05-0.1% trypsin for 5-10 min at 37°C. For E6.0 and E6.6 embryos, epiblast cells were mechanically dissociated to obtain small cell clusters (<10 cells), which were then treated with TryPLE for 5 min at 37°C, and singularized by gentle mechanical dissociation. Enzymatic activities were stopped by adding 10% fetal bovine serum (Gibco 11563397). Cell suspensions were run through a 50μm filter to remove any remaining cell clumps and were loaded on a Chromium controller.

### RNA extraction and bulk RNA sequencing

Total RNA was isolated from batches of embryos, ICM or TE (n = 20) using PicoPur Arcturus (Excilone, France) with a DNase I (Qiagen, Germany) treatment as recommended by the supplier. For E6.0, E6.3, and E6.6 stages, RNAs were extracted from single embryo samples. Three nanograms of total RNA were used for amplification using the SMART-Seq V4 Ultra Low Input RNA kit (Clontech) according to the manufacturer’s recommendations (10 PCR cycles were performed). cDNA quality was assessed on an Agilent Bioanalyzer 2100, using an Agilent High Sensitivity DNA Kit. Libraries were prepared from 0.15 ng cDNA using the Nextera XT Illumina library preparation kit. Libraries were pooled in equimolar proportions and sequenced (Paired-end 50–34 bp) on an Illumina NextSeq500 instrument, using a NextSeq 500 High Output 75 cycles kit. Demultiplexing was performed (bcl2fastq2V2.2.18.12) and adapters were trimmed with Cutadapt1.15, so that only reads longer than 10 bp were kept. Number of reads ranged from 52 to 137 million. Reads were mapped to the rabbit genome (Oryctolagus cuniculus 2.0). 81.8 to 85.7% (depending on samples) of the pair fragments could be aligned; 70.3 to 78% of these fragments passed the mapping filter; 56.4 to 64.7% of them were assigned to a gene.

### Single-cell RNA-library construction and sequencing

Cell suspensions were loaded on a Chromium controller (10x Genomics, Pleasanton, CA, USA) to generate single-cell Gel Beads-in-Emulsion (GEMs). Single-cell RNAseq libraries were prepared using Chromium Single cell 3’RNA Gel Bead and Library Kit (P/N 1000075, 1000153, 10x Genomics). GEM-RT was performed in a C1000 Touch Thermal cycler with 96-Deep Well Reaction Module (Bio-Rad; P/N 1851197): 53°C for 45 min, 85°C for 5 min; held at 4°C. After RT, GEMs were broken and the single-strand cDNA was cleaned up with DynaBeads MyOne Silane Beads (Thermo Fisher Scientific; P/N 37002D). cDNA was amplified using the C1000 Touch Thermal cycler with 96-DeepWell Reaction Module: 98°C for 3 min; 11 cycle of 98°C for 15 s, 63°C for 20 s, and 72°C for 1 min; 72°C for 1 min; held at 4°C. Amplified cDNA product was cleaned up with the SPRIselect beads (SPRI P/N B23318). Indexed sequencing libraries were constructed with SPRIselect following these steps: (1) Fragmentation end repair and A-tailing and size selection; (2) adapter ligation and cleanup; (3) sample index PCR and size selection. The barcode sequencing libraries were quantified by quantitative PCR (KAPA Biosystems Library Quantification Kit for Illumina platforms P/N KK4824). Sequencing libraries were loaded at 18 pM on an Illumina HiSeq2500 using the following read length: 28 bp Read1, 8 bp I7 Index, 91 bp Read2.

### Bioinformatics analysis

For bulk RNAseq, reads were mapped to the rabbit genome (Oryctolagus cuniculus 2.0) using the splice junction mapper TopHat (version 2.0.14) associated with the short-read aligner Bowtie2 (version 2.1.0). Finally, FeatureCounts (version 1.4.5) was then used to establish a gene count table. Hierarchical clustering was computed using hclust R package with Euclidean distance metric and Ward linkage. Principal Component analysis (PCA) was made using FactoMineR (Lê et al., 2008). Data normalization and single-gene level analysis of differentially expression genes were performed using the DESeq2 (Anders and Huber, 2010). Differences were considered significant when adjusted P values (Benjamini-Hochberg) < 0.01, and absolute fold change ≥ 2. Log2 transformed read counts (Rlog) (after normalization by library size) were obtained with DEseq2 package and used for heatmaps.

For single-cell RNAseq, data were analysed using the R software (version 3.6.3) and RStudio Desktop integrated development environment (IDE) for R (version 1.4, Open Source Edition). Since the rabbit genome annotation is not complete, some genes were annotated manually using ENSG identity. The Ensembl release 104 (Howe et al., 2021), the National Center for Biotechnology Information (NCBI database) (Schoch et al., 2020), and the g:Profiler online software (Raudvere et al., 2019) were used to convert ENSG annotations into Gene symbols. The Seurat package (version 3.1.2) was used to filter, normalize and analyse the data as described (Stuart et al., 2019). Briefly, the data were filtered by eliminating all cells with mitochondrial genome expression above 0.7%, and selecting cells in which the total number of expressed genes is between 300 and 2500. A fraction of the cells expressed mitochondrial genes at very low levels. We hypothesized that these were nuclei that had been stripped of their cytoplasm during embryo dissection, single-cell dissociation, or single-cell capture in 10x droplets, and were removed from the analyses. In a previous study by Pijuan-Sala and collaborators, similar artefacts were observed and the corresponding events were eliminated (Pijuan-Sala et al., 2019). For the remaining cells, the novelty scores were above 0.80 (**Fig S1D**) as expected for good quality cells.

Expression levels were then log-normalized and the 3000 most variable genes were identified. Principal component analysis (PCA) and uniform manifold approximation and projection (UMAP) dimension reductions were applied to the dataset to identify cell clusters. These clusters were identified using graph-based clustering algorithm on the first 20 components and annotated according to expression of lineage-specific genes and embryonic stage. Differential expression analysis was performed and genes with a log fold-change > 0.25 or < −0.25 were considered significantly differentially expressed. The same workflow was applied for published data.

Dataset integration was done by retrieving the data from Nakamura et al., 2016 on mouse (GSE63266) and cynomolgus monkey embryos (GSE74767) and merging them with our own 10x dataset. The original annotations for the mouse and cynomolgus monkey cells were used. All three datasets were first processed using Seurat (Stuart et al., 2019) as previously described. After normalization, the data were integrated using 4000 genes and 20 dimensions with the Find Integration Anchors function. PCA and UMAP were then used on the merged dataset in order to localize the different embryonic lineages.

Temporal trajectories were created using the following additional packages: monocle (version 2.12.0), cellranger (version 1.1.0) and viridislite (version 0.4.0). Data analysis was performed as described (Trapnell et al., 2014). Briefly, differential analysis was performed on cells isolated and annotated using Seurat package, and the top 1000 most significantly differentially expressed genes were used to order the samples in pseudo-time. Stage E3.0 was set as the starting point.

Statistical enrichment analysis of Gene Ontology (GO) and Kyoto Encyclopedia of Genes and Genomes (KEGG) terms was performed using the g:Profiler online software (Raudvere et al., 2019). Cytoscape software (Shannon et al., 2003; Smoot et al., 2011) and enrichment map plug-ins were used to build and visualize networks. The node size is scaled by the number of genes contributing to each term; edges width is proportional to the overlap between each gene set and each node is coloured by their enrichment score.

### Immunofluorescence

Embryos were fixed in 4% paraformaldehyde (PFA) for 20 min at room temperature. After three washes in phosphate-buffered saline (PBS), embryos were permeabilized in PBS-0.5% TritonX100 for 30 min. For anti-5’hydroxymethylcytosine and 5’methylcytosine antibodies, embryos were permeabilized in PBS-0.5% Triton for 15 min, washed in PBS for 20 min, then incubated in 2M HCl for 30 min. Embryos were subsequently incubated in 100mM Tris-HCl pH 9 for 10 min, washed in PBS for 15 min, and incubated in blocking solution (PBS supplemented with 5% bovine serum albumin) for 1 h at room temperature. Embryos were then incubated with primary antibodies diluted in blocking solution overnight at 4°C (**Table S1**). After two washes (2 x 15 min) in PBS, embryos were incubated in secondary antibodies diluted in blocking solution at a dilution of 1:100 for 1 h at room temperature. Finally, embryos were transferred through several washes of PBS before staining DNA with 4’,6-diamidino-2-phenylindole (DAPI; 0.5μg/mL) or propidium iodide (PI; 1μg/mL) for 10 min at room temperature. Embryos were analysed by confocal imaging (DM 6000 CS SP5; Leica). Acquisitions were performed using an oil immersion objective (40x/1.25 0.75, PL APO HCX; Leica).

### Detection of ROS, glycolysis, and mitochondrial membrane potential

After gentle removal of the mucin coat with protease from Streptomyces griseus (5mg/mL), embryos were incubated in pre-warmed pre-equilibrated RDH medium containing 4μM Hoechst 33342 for DNA staining and either 5μM CellROX Deep Red reagent, 5μM Image-iT Red Hypoxia reagent or 50 nM TMRE (all from ThermoFisher Scientific) for 30-45 min at 38°C. For glucose uptake assays, embryos were incubated for 30 min in the glucose uptake mix, according to the manufacturer’s instructions (Abcam ab204702). Embryos were then collected and washed twice with the analysis buffer. After staining, embryos were transferred to new drops of RDH before mounting in M2 medium (Sigma) containing 20% Optiprep (Stem Cell Technologies) and ProLong Live antifade reagent for live cell imaging (1/100 dilution, Thermofisher). Embryos were then imaged on a Leica SP5 confocal microscope with a x25 water immersion objective.

## Acknowledgments

We thank the members of UE1298 SAAJ Science de l’Animal et de l’Aliment, responsible for INRAE Jouy-en-Josas rabbit facility; as well as the High-throughput sequencing facility of I2BC.

## Footnotes

### Author contributions

Conceptualization, N.B, P.S., M.A., and V.D.; Methodology, N.B., M.A., W.B., M.G., and V.D.; Validation, W.B., N.B., and V.D.; Formal Analysis, W.B., L.J., C.A.; Investigation, W.B., C.A., I.A., A.M., N.D., N.P., S.C., Y.J., M.P., and D.S.; Resources, T.J.; Data Curation, W.B. and L.J.; Writing - Original Draft, P.S. and N.B.; Writing – Review and Editing, W.B., M.A., and V.D.; Visualization, W.B. and N.B; Supervision: N.B.; Project Administration, M.A., N.B, and V.D.; Funding Acquisition, P.S. and V.D.

### Data availability

The bulk RNAseq datasets generated in this study are deposited in NCBI SRA with the accession numbers: PRJNA529333 for TE_96h libraries and PRJNA743177 for whole embryo and tissue data. For the 10x single-cell RNAseq datasets, all raw read sequence data, count matrices for each stage and count matrix of the aggregated datasets from each stage generated in this study are deposited in NCBI GEO with the accession number: GSE180048.

## References

Aksoy, I., Rognard, C., Moulin, A., Marcy, G., Masfaraud, E., Wianny, F., Cortay, V., Bellemin-Menard, A., Doerflinger, N., Dirheimer, M., et al. (2021). Apoptosis, G1 Phase Stall, and Premature Differentiation Account for Low Chimeric Competence of Human and Rhesus Monkey Naive Pluripotent Stem Cells. Stem cell reports 16, 56–74.

Anders, S. and Huber, W. (2010). Differential expression analysis for sequence count data. Genome Biol 11, R106.

Argelaguet, R., Clark, S. J., Mohammed, H., Stapel, L. C., Krueger, C., Kapourani, C. A., Imaz-Rosshandler, I., Lohoff, T., Xiang, Y., Hanna, C. W., et al. (2019). Multi-omics profiling of mouse gastrulation at single-cell resolution. Nature 576, 487–491.

Bayerl, J., Ayyash, M., Shani, T., Manor, Y. S., Gafni, O., Massarwa, R., Kalma, Y., Aguilera-Castrejon, A., Zerbib, M., Amir, H., et al. (2021). Principles of signaling pathway modulation for enhancing human naive pluripotency induction. Cell Stem Cell 28, 1–17.

Blakeley, P., Fogarty, N. M., Del Valle, I., Wamaitha, S. E., Hu, T. X., Elder, K., Snell, P., Christie, L., Robson, P. and Niakan, K. K. (2015). Defining the three cell lineages of the human blastocyst by single-cell RNA-seq. Development 142, 3151–3165.

Boroviak, T., Loos, R., Lombard, P., Okahara, J., Behr, R., Sasaki, E., Nichols, J., Smith, A. and Bertone, P. (2015). Lineage-Specific Profiling Delineates the Emergence and Progression of Naive Pluripotency in Mammalian Embryogenesis. Dev Cell 35, 366–382.

Boroviak, T., Stirparo, G. G., Dietmann, S., Hernando-Herraez, I., Mohammed, H., Reik, W., Smith, A., Sasaki, E., Nichols, J. and Bertone, P. (2018). Single cell transcriptome analysis of human, marmoset and mouse embryos reveals common and divergent features of preimplantation development. Development 145, dev167833.

Bredenkamp, N., Stirparo, G. G., Nichols, J., Smith, A. and Guo, G. (2019). The Cell-Surface Marker Sushi Containing Domain 2 Facilitates Establishment of Human Naive Pluripotent Stem Cells. Stem cell reports 12, 1212–1222.

Brons, I. G., Smithers, L. E., Trotter, M. W., Rugg-Gunn, P., Sun, B., Chuva de Sousa Lopes, S. M., Howlett, S. K., Clarkson, A., Ahrlund-Richter, L., Pedersen, R. A., et al. (2007). Derivation of pluripotent epiblast stem cells from mammalian embryos. Nature 448, 191–195.

Canon, E., Jouneau, L., Blachere, T., Peynot, N., Daniel, N., Boulanger, L., Maulny, L., Archilla, C., Voisin, S., Jouneau, A., et al. (2018). Progressive methylation of POU5F1 regulatory regions during blastocyst development. Reproduction 156, 145–161.

Carbognin, E., Betto, R. M., Soriano, M. E., Smith, A. G. and Martello, G. (2016). Stat3 promotes mitochondrial transcription and oxidative respiration during maintenance and induction of naive pluripotency. EMBO J 35, 618–634.

Cassar, P. A., Carpenedo, R. L., Samavarchi-Tehrani, P., Olsen, J. B., Park, C. J., Chang, W. Y., Chen, Z., Choey, C., Delaney, S., Guo, H., et al. (2015). Integrative genomics positions MKRN1 as a novel ribonucleoprotein within the embryonic stem cell gene regulatory network. EMBO Rep 16, 1334–1357.

Chaumeil, J., Waters, P. D., Koina, E., Gilbert, C., Robinson, T. J. and Graves, J. A. (2011). Evolution from XIST-independent to XIST-controlled X-chromosome inactivation: epigenetic modifications in distantly related mammals. PLoS ONE 6, e19040.

Chazaud, C. and Yamanaka, Y. (2016). Lineage specification in the mouse preimplantation embryo. Development 143, 1063–1074.

Corsinotti, A., Wong, F. C., Tatar, T., Szczerbinska, I., Halbritter, F., Colby, D., Gogolok, S., Pantier, R., Liggat, K., Mirfazeli, E. S., et al. (2017). Distinct SoxB1 networks are required for naive and primed pluripotency. eLife 6, e27746.

Davidson, K. C., Mason, E. A. and Pera, M. F. (2015). The pluripotent state in mouse and human. Development 142, 3090–3099.

Devika, A. S., Wruck, W., Adjaye, J. and Sudheer, S. (2019). The quest for pluripotency: a comparative analysis across mammalian species. Reproduction 158, R97–R111.

Dunn, S. J., Martello, G., Yordanov, B., Emmott, S. and Smith, A. G. (2014). Defining an essential transcription factor program for naive pluripotency. Science 344, 1156–1160.

Fischer, B., Chavatte-Palmer, P., Viebahn, C., Navarrete Santos, A. and Duranthon, V. (2012). Rabbit as a reproductive model for human health. Reproduction 144, 1–10.

Guo, G., Huss, M., Tong, G. Q., Wang, C., Li Sun, L., Clarke, N. D. and Robson, P. (2010). Resolution of cell fate decisions revealed by single-cell gene expression analysis from zygote to blastocyst. Dev Cell 18, 675–685.

Habibi, E., Brinkman, A. B., Arand, J., Kroeze, L. I., Kerstens, H. H., Matarese, F., Lepikhov, K., Gut, M., Brun-Heath, I., Hubner, N. C., et al. (2013). Whole-genome bisulfite sequencing of two distinct interconvertible DNA methylomes of mouse embryonic stem cells. Cell Stem Cell 13, 360–369.

Hackett, J. A., Sengupta, R., Zylicz, J. J., Murakami, K., Lee, C., Down, T. A. and Surani, M. A. (2013). Germline DNA demethylation dynamics and imprint erasure through 5-hydroxymethylcytosine. Science 339, 448–452.

Hassoun, R., Puschel, B. and Viebahn, C. (2009a). Sox17 Expression Patterns during Gastrulation and Early Neurulation in the Rabbit Suggest Two Sources of Endoderm Formation. Cells Tissues Organs 191, 68–83.

Hassoun, R., Schwartz, P., Feistel, K., Blum, M. and Viebahn, C. (2009b). Axial differentiation and early gastrulation stages of the pig embryo. Differentiation 78, 301–311.

Hayashi, K., de Sousa Lopes, S. M. and Surani, M. A. (2007). Germ cell specification in mice. Science 316, 394–396.

Hayashi, K., Lopes, S. M., Tang, F. and Surani, M. A. (2008). Dynamic equilibrium and heterogeneity of mouse pluripotent stem cells with distinct functional and epigenetic states. Cell Stem Cell 3, 391–401.

Hopf, C., Viebahn, C. and Puschel, B. (2011). BMP signals and the transcriptional repressor BLIMP1 during germline segregation in the mammalian embryo. Development genes and evolution 221, 209–223.

Houghton, F. D. (2006). Energy metabolism of the inner cell mass and trophectoderm of the mouse blastocyst. Differentiation 74, 11–18.

Howe, K. L., Achuthan, P., Allen, J., Allen, J., Alvarez-Jarreta, J., Amode, M. R., Armean, I. M., Azov, A. G., Bennett, R., Bhai, J., et al. (2021). Ensembl 2021. Nucleic Acids Res 49, D884–D891.

Kalkan, T., Bornelov, S., Mulas, C., Diamanti, E., Lohoff, T., Ralser, M., Middelkamp, S., Lombard, P., Nichols, J. and Smith, A. (2019). Complementary Activity of ETV5, RBPJ, and TCF3 Drives Formative Transition from Naive Pluripotency. Cell Stem Cell 24, 785–801 e787.

Kinoshita, M., Barber, M., Mansfield, W., Cui, Y., Spindlow, D., Stirparo, G. G., Dietmann, S., Nichols, J. and Smith, A. (2020). Capture of Mouse and Human Stem Cells with Features of Formative Pluripotency. Cell Stem Cell 28, 453–471 e458.

Kobayashi, T., Castillo-Venzor, A., Penfold, C. A., Morgan, M., Mizuno, N., Tang, W. W.C., Osada, Y., Hirao, M., Yoshida, F., Sato, H., et al. (2021). Tracing the emergence of primordial germ cells from bilaminar disc rabbit embryos and pluripotent stem cells. Cell reports 37, 109812.

Kong, Q., Yang, X., Zhang, H., Liu, S., Zhao, J., Zhang, J., Weng, X., Jin, J. and Liu, Z. (2020). Lineage specification and pluripotency revealed by transcriptome analysis from oocyte to blastocyst in pig. FASEB J 34, 691–705.

Lê, S., Josse, J. and Husson, F. (2008). FactoMineR: an R package for multivariate analysis. J Stat Softw 25, 1–18.

Leandri, R. D., Archilla, C., Bui, L. C., Peynot, N., Liu, Z., Cabau, C., Chastellier, A., Renard, J. P. and Duranthon, V. (2009). Revealing the dynamics of gene expression during embryonic genome activation and first differentiation in the rabbit embryo with a dedicated array screening. Physiol Genomics 36, 98–113.

Lee, E. W., Lee, M. S., Camus, S., Ghim, J., Yang, M. R., Oh, W., Ha, N. C., Lane, D. P. and Song, J. (2009). Differential regulation of p53 and p21 by MKRN1 E3 ligase controls cell cycle arrest and apoptosis. EMBO J 28, 2100–2113.

Leitch, H. G., McEwen, K. R., Turp, A., Encheva, V., Carroll, T., Grabole, N., Mansfield, W., Nashun, B., Knezovich, J. G., Smith, A., et al. (2013). Naive pluripotency is associated with global DNA hypomethylation. Nature structural & molecular biology 20, 311–316

Leitch, H. G., Surani, M. A. and Hajkova, P. (2016). DNA (De)Methylation: The Passive Route to Naivety? Trends Genet 32, 592–595.

Levine, A. J. and Brivanlou, A. H. (2006). GDF3, a BMP inhibitor, regulates cell fate in stem cells and early embryos. Development 133, 209–216.

Liu, D., Wang, X., He, D., Sun, C., He, X., Yan, L., Li, Y., Han, J. J. and Zheng, P. (2018). Single cell RNA-sequencing reveals the existence of naive and primed pluripotency in pre-implantation rhesus monkey embryos. Genome Res 28, 1481–1493.

Mason, M.J., Fan, G., Plath, K., Zhou, Q. and Horvath, S. (2009). Signed weighted gene co-expression network analysis of transcriptional regulation in murine embryonic stem cells. BMC Genomics 10, 327.

Meistermann, D., Bruneau, A., Loubersac, S., Reignier, A., Firmin, J., Francois-Campion, V., Kilens, S., Lelievre, Y., Lammers, J., Feyeux, M., et al. (2021). Integrated pseudotime analysis of human pre-implantation embryo single-cell transcriptomes reveals the dynamics of lineage specification. Cell Stem Cell 28, 1625–1640 e1626.

Messmer, T., von Meyenn, F., Savino, A., Santos, F., Mohammed, H., Lun, A. T. L., Marioni, J. C. and Reik, W. (2019). Transcriptional Heterogeneity in Naive and Primed Human Pluripotent Stem Cells at Single-Cell Resolution. Cell reports 26, 815–824 e814.

Mohammed, H., Hernando-Herraez, I., Savino, A., Scialdone, A., Macaulay, I., Mulas, C., Chandra, T., Voet, T., Dean, W., Nichols, J., et al. (2017). Single-Cell Landscape of Transcriptional Heterogeneity and Cell Fate Decisions during Mouse Early Gastrulation. Cell reports 20, 1215–1228.

Nakamura, T., Fujiwara, K., Saitou, M. and Tsukiyama, T. (2021). Non-human primates as a model for human development. Stem cell reports 16, 1093–1103.

Nakamura, T., Okamoto, I., Sasaki, K., Yabuta, Y., Iwatani, C., Tsuchiya, H., Seita, Y., Nakamura, S., Yamamoto, T. and Saitou, M. (2016). A developmental coordinate of pluripotency among mice, monkeys and humans. Nature 537, 57–62.

Nowotschin, S., Setty, M., Kuo, Y. Y., Liu, V., Garg, V., Sharma, R., Simon, C. S., Saiz, N., Gardner, R., Boutet, S. C., et al. (2019). The emergent landscape of the mouse gut endoderm at single-cell resolution. Nature 569, 361–367.

Okamoto, I., Patrat, C., Thepot, D., Peynot, N., Fauque, P., Daniel, N., Diabangouaya, P., Wolf, J. P., Renard, J. P., Duranthon, V., et al. (2011). Eutherian mammals use diverse strategies to initiate X-chromosome inactivation during development. Nature 472, 370–374.

Osteil, P., Moulin, A., Santamaria, C., Joly, T., Jouneau, L., Aubry, M., Tapponnier, Y., Archilla, C., Schmaltz-Panneau, B., Lecardonnel, J., et al. (2016). A Panel of Embryonic Stem Cell Lines Reveals the Variety and Dynamic of Pluripotent States in Rabbits. Stem cell reports 7, 383–398.

Peng, G., Suo, S., Chen, J., Chen, W., Liu, C., Yu, F., Wang, R., Chen, S., Sun, N., Cui, G., et al. (2016). Spatial Transcriptome for the Molecular Annotation of Lineage Fates and Cell Identity in Mid-gastrula Mouse Embryo. Dev Cell 36, 681–697.

Petropoulos, S., Edsgard, D., Reinius, B., Deng, Q., Panula, S. P., Codeluppi, S., Plaza Reyes, A., Linnarsson, S., Sandberg, R. and Lanner, F. (2016). Single-Cell RNA-Seq Reveals Lineage and X Chromosome Dynamics in Human Preimplantation Embryos. Cell 165, 1012–1026.

Pierre, A., Gautier, M., Callebaut, I., Bontoux, M., Jeanpierre, E., Pontarotti, P. and Monget, P. (2007). Atypical structure and phylogenomic evolution of the new eutherian oocyte- and embryo-expressed KHDC1/DPPA5/ECAT1/OOEP gene family. Genomics 90, 583–594.

Pijuan-Sala, B., Griffiths, J. A., Guibentif, C., Hiscock, T. W., Jawaid, W., Calero-Nieto, F. J., Mulas, C., Ibarra-Soria, X., Tyser, R. C. V., Ho, D. L. L., et al. (2019). A single-cell molecular map of mouse gastrulation and early organogenesis. Nature 566, 490–495.

Piliszek, A., Madeja, Z. E. and Plusa, B. (2017). Suppression of ERK signalling abolishes primitive endoderm formation but does not promote pluripotency in rabbit embryo. Development 144, 3719–3730.

Puschel, B., Daniel, N., Bitzer, E., Blum, M., Renard, J. P. and Viebahn, C. (2010). The rabbit (Oryctolagus cuniculus): a model for mammalian reproduction and early embryology. Cold Spring Harbor protocols 2010, pdb emo139.

Qian, X., Kim, J. K., Tong, W., Villa-Diaz, L. G. and Krebsbach, P. H. (2016). DPPA5 Supports Pluripotency and Reprogramming by Regulating NANOG Turnover. Stem Cells 34, 588–600.

Raudvere, U., Kolberg, L., Kuzmin, I., Arak, T., Adler, P., Peterson, H. and Vilo, J. (2019). g:Profiler: a web server for functional enrichment analysis and conversions of gene lists (2019 update). Nucleic Acids Res 47, W191–W198.

Sachs, P., Ding, D., Bergmaier, P., Lamp, B., Schlagheck, C., Finkernagel, F., Nist, A., Stiewe, T. and Mermoud, J. E. (2019). SMARCAD1 ATPase activity is required to silence endogenous retroviruses in embryonic stem cells. Nat Commun 10, 1335.

Savatier, P., Osteil, P. and Tam, P. P. (2017). Pluripotency of embryo-derived stem cells from rodents, lagomorphs, and primates: Slippery slope, terrace and cliff. Stem Cell Res 19, 104–112.

Schmaltz-Panneau, B., Jouneau, L., Osteil, P., Tapponnier, Y., Afanassieff, M., Moroldo, M., Jouneau, A., Daniel, N., Archilla, C., Savatier, P., et al. (2014). Contrasting transcriptome landscapes of rabbit pluripotent stem cells in vitro and in vivo. Anim Reprod Sci 149, 67–79.

Schoch, C. L., Ciufo, S., Domrachev, M., Hotton, C. L., Kannan, S., Khovanskaya, R., Leipe, D., McVeigh, R., O’Neill, K., Robbertse, B., et al. (2020). NCBI Taxonomy: a comprehensive update on curation, resources and tools. Database (Oxford) 2020.

Shahbazi, M. N. (2020). Mechanisms of hu.man embryo development: from cell fate to tissue shape and back. Development 147, dev190629

Shannon, P., Markiel, A., Ozier, O., Baliga, N. S., Wang, J. T., Ramage, D., Amin, N., Schwikowski, B. and Ideker, T. (2003). Cytoscape: a software environment for integrated models of biomolecular interaction networks. Genome Res 13, 2498–2504.

Smith, A. (2017). Formative pluripotency: the executive phase in a developmental continuum. Development 144, 365–373.

Smoot, M. E., Ono, K., Ruscheinski, J., Wang, P. L. and Ideker, T. (2011). Cytoscape 2.8: new features for data integration and network visualization. Bioinformatics 27, 431–432.

Sone, M., Morone, N., Nakamura, T., Tanaka, A., Okita, K., Woltjen, K., Nakagawa, M., Heuser, J. E., Yamada, Y., Yamanaka, S., et al. (2017). Hybrid Cellular Metabolism Coordinated by Zic3 and Esrrb Synergistically Enhances Induction of Naive Pluripotency. Cell metabolism 25, 1103–1117 e1106.

Stirparo, G. G., Boroviak, T., Guo, G., Nichols, J., Smith, A. and Bertone, P. (2018). Integrated analysis of single-cell embryo data yields a unified transcriptome signature for the human preimplantation epiblast. Development 145, dev158501.

Stuart, T., Butler, A., Hoffman, P., Hafemeister, C., Papalexi, E., Mauck, W. M., 3rd, Hao, Y., Stoeckius, M., Smibert, P. and Satija, R. (2019). Comprehensive Integration of Single-Cell Data. Cell 177, 1888–1902 e1821.

Takahashi, S., Kobayashi, S. and Hiratani, I. (2017). Epigenetic differences between naive and primed pluripotent stem cells. Cell Mol Life Sci 75, 1191–1203.

Tanaka, T. S., Lopez de Silanes, I., Sharova, L. V., Akutsu, H., Yoshikawa, T., Amano, H., Yamanaka, S., Gorospe, M. and Ko, M. S. (2006). Esg1, expressed exclusively in preimplantation embryos, germline, and embryonic stem cells, is a putative RNA-binding protein with broad RNA targets. Dev Growth Differ 48, 381–390.

Tang, F., Barbacioru, C., Bao, S., Lee, C., Nordman, E., Wang, X., Lao, K. and Surani, M. A. (2010). Tracing the derivation of embryonic stem cells from the inner cell mass by single-cell RNA-Seq analysis. Cell Stem Cell 6, 468–478.

Tapponnier, Y., Afanassieff, M., Aksoy, I., Aubry, M., Moulin, A., Medjani, L., Bouchereau, W., Mayere, C., Osteil, P., Nurse-Francis, J., et al. (2017). Reprogramming of rabbit induced pluripotent stem cells toward epiblast and chimeric competency using Kruppel-like factors. Stem Cell Res 24, 106–117.

Teixeira, M., Commin, L., Gavin-Plagne, L., Bruyere, P., Buff, S. and Joly, T. (2018). Rapid cooling of rabbit embryos in a synthetic medium. Cryobiology 85, 113–119.

Tesar, P. J., Chenoweth, J. G., Brook, F. A., Davies, T. J., Evans, E. P., Mack, D. L., Gardner, R. L. and McKay, R. D. (2007). New cell lines from mouse epiblast share defining features with human embryonic stem cells. Nature 448, 196–199.

Trapnell, C., Cacchiarelli, D., Grimsby, J., Pokharel, P., Li, S., Morse, M., Lennon, N. J., Livak, K. J., Mikkelsen, T. S. and Rinn, J. L. (2014). The dynamics and regulators of cell fate decisions are revealed by pseudotemporal ordering of single cells. Nat Biotechnol 32, 381–386.

Tripathi, P., Dakle, P., Mehravar, M., Pandey, V. K., Bullen, M. J., Zhang, Z., Hathiwala, D., Kerenyi, M., Woo, A., Ghamari, A., et al. (2021). Histone demethylome map reveals combinatorial gene regulatory functions in embryonic stem cells. bioRxiv, 2020.2008.2027.269514.

Viebahn, C., Stortz, C., Mitchell, S. A. and Blum, M. (2002). Low proliferative and high migratory activity in the area of Brachyury expressing mesoderm progenitor cells in the gastrulating rabbit embryo. Development 129, 2355–2365.

Wen, J., Zeng, Y., Fang, Z., Gu, J., Ge, L., Tang, F., Qu, Z., Hu, J., Cui, Y., Zhang, K., et al. (2017). Single-cell analysis reveals lineage segregation in early post-implantation mouse embryos. J Biol Chem 292, 9840–9854.

Wianny, F., Real, F. X., Mummery, C. L., Van Rooijen, M., Lahti, J., Samarut, J. and Savatier, P. (1998). G1-phase regulators, cyclin D1, cyclin D2, and cyclin D3: up-regulation at gastrulation and dynamic expression during neurulation. Dev Dyn 212, 49–62.

Xiao, S., Lu, J., Sridhar, B., Cao, X., Yu, P., Zhao, T., Chen, C. C., McDee, D., Sloofman, L., Wang, Y., et al. (2017). SMARCAD1 Contributes to the Regulation of Naive Pluripotency by Interacting with Histone Citrullination. Cell reports 18, 3117–3128.

Zhang, Y., Xiang, Y., Yin, Q., Du, Z., Peng, X., Wang, Q., Fidalgo, M., Xia, W., Li, Y., Zhao, Z. A., et al. (2018). Dynamic epigenomic landscapes during early lineage specification in mouse embryos. Nat Genet 50, 96–105.

Zhao, X. M., Cui, L. S., Hao, H. S., Wang, H. Y., Zhao, S. J., Du, W. H., Wang, D., Liu, Y. and Zhu, H. B. (2016). Transcriptome analyses of inner cell mass and trophectoderm cells isolated by magnetic-activated cell sorting from bovine blastocysts using single cell RNA-seq. Reprod Domest Anim 51, 726–735.

Zhou, W., Choi, M., Margineantu, D., Margaretha, L., Hesson, J., Cavanaugh, C., Blau, C. A., Horwitz, M. S., Hockenbery, D., Ware, C., et al. (2012). HIF1alpha induced switch from bivalent to exclusively glycolytic metabolism during ESC-to-EpiSC/hESC transition. EMBO J 31, 2103–2116.

Zhu, P., Guo, H., Ren, Y., Hou, Y., Dong, J., Li, R., Lian, Y., Fan, X., Hu, B., Gao, Y., et al. (2018). Single-cell DNA methylome sequencing of human preimplantation embryos. Nat Genet 50, 12–19.

