## Supplementary figures for "Major transcriptomic, epigenetic and metabolic changes underly the pluripotency continuum in rabbit preimplantation embryos"

### Figure S1: Raw data processing and quality controls.

A

| Embryo stages | E2.7 | E3.5 | | E4.0 | | E6.0 | | | E6.3 | E6.6 | |
|---|---|---|---|---|---|---|---|---|---|---|---|
| Embryo tissues | whole | ICM | TE | ICM | TE | EPI | PE | TE | EPI_ant | EPI_ant | TE |
| Number ofbiological replicates | 3 | 3 | 3 | 3 | 3 | 6 | 6 | 6 | 6 | 6 | 6 |
| Number of embryos/replicate | 20 | 20 | 20 | 20 | 20 | 1 | 1 | 1 | 1 | 1 | 1 |
| Reads (in million) | 54.3-137.8 | 64.0-79.7 | 66.5-81.8 | 70.2-86.0 | 67.4-73.1 | 64.5-85.0 | 70.9-90.0 | 53.8-90.0 | 52.9-84.8 | 54.3-91.4 | 57.4-93.3 |
| % of aligned pairs | 82.8-85.3 | 81.8-85.7 | 84.6-84.9 | 83.6-84.8 | 83.2-84.7 | 84.0-84.8 | 83.2-85.0 | 84.4-84.9 | 83.3-85.3 | 83.5-85.4 | 84.7-85.5 |
| % of reads assigned to gene | 60.6-62.7 | 58.1-61.6 | 62.0-62.5 | 60.2-60.4 | 61.3-63.9 | 59.0-59.8 | 58.3-61.0 | 62.3-63.3 | 56.7-59.5 | 56.4-59.7 | 62.4-64.7 |

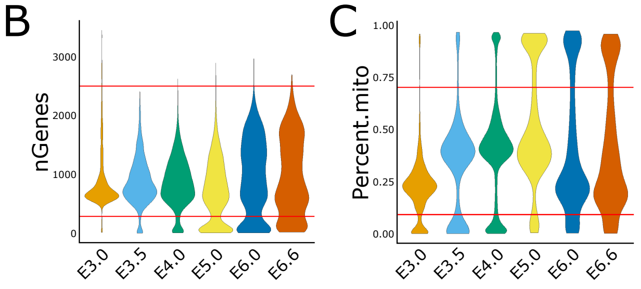


D

| | *E3.0* | *E3.5* | *E4.0* | *E5.0* | *E6.0* | *E6.6* |
|---|---|---|---|---|---|---|
| Dissociated embryos | 132 | 166 | 76 | 90 | 28 | 37 |
| Sequenced cells | 1 294 | 2 853 | 5 227 | 5 274 | 3 453 | 2 791 |
| Analysed cells | 1 051 | 1 943 | 3 795 | 3 251 | 2 119 | 1 783 |
| Mean reads/cell | 56 103 | 14 705 | 15 412 | 22 139 | 15 293 | 23 314 |
| Median genes/cell | 1 096 | 877 | 944 | 925 | 948 | 1 087 |
| Median UMI/cell | 4 297 | 3 399 | 3 941 | 4 852 | 3 407 | 4 681 |
| Log10 genes /Log10 UMI | 0.84 | 0.83 | 0.83 | 0.80 | 0.84 | 0.83 |

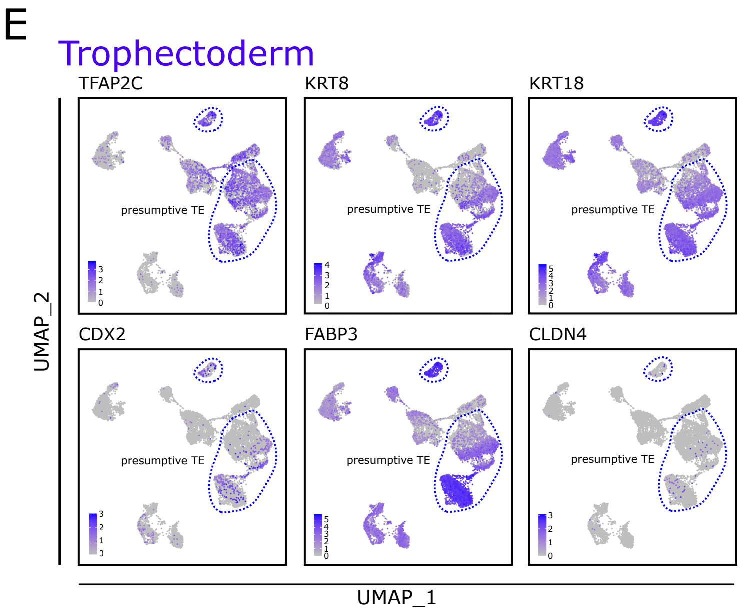


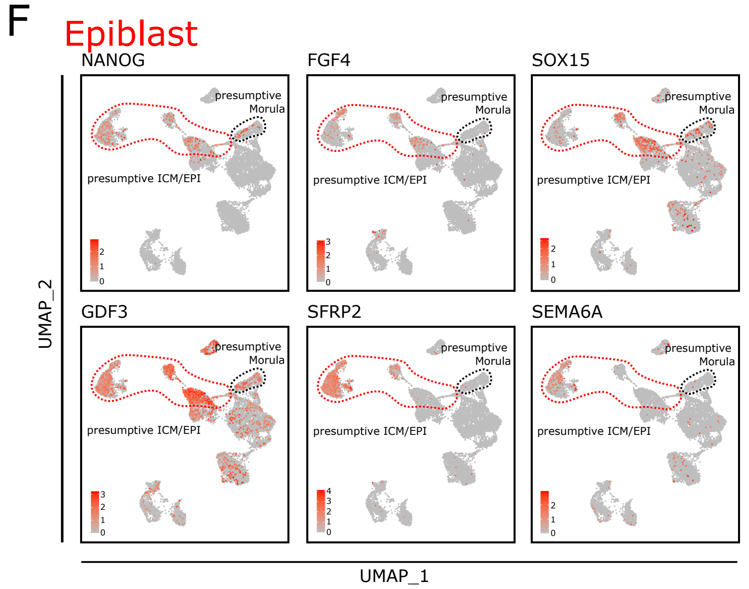


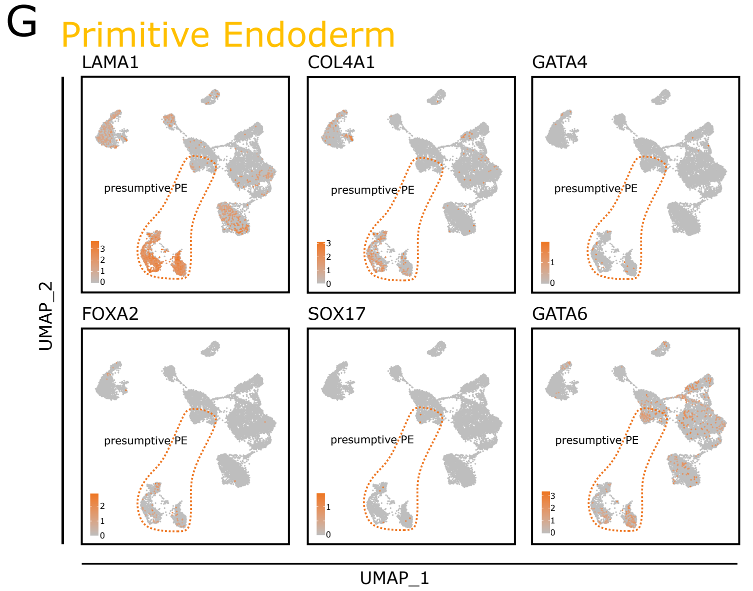


**Figure S1: Raw data processing and quality controls.** (**A**) Sample description (bulk RNAseq) for each developmental stage (E2.7 to E6.6) with numbers of biological replicates, number of embryos per replicate, numbers of reads per library (minimal and maximal values), percentages of aligned pairs (minimal and maximal values), and percentages of reads assigned to a gene (minimal and maximal values). (**B**) Violin plot representation of the number of genes detected in the 10X Genomics single-cell RNAseq dataset for each developmental stage analysed (E3.0 to E6.6); red lines indicate the thresholds used for sample filtering with retention of cells only displaying expression of more than 300 genes and less than 2500 genes. (**C**) Violin plot representation of the percentage of mitochondrial genes detected in the 10X Genomics single-cell RNAseq dataset for each developmental stage analysed (E3.0 to E6.6). Red line indicates the threshold applied to samples filtering with retention of cells only displaying more than 0.1% and less than 0.7% of mitochondrial genome. (**D**) Sample description (10X Genomics single-cell RNAseq) for each developmental stage (E3.0 to E6.6) with numbers of dissociated embryos, analysed cells, cells with sequenced genomes, means of reads, median of genes detected per cell, unique molecular identifier (UMI) per cell and novelty score (Log10 nGenes/Log10 nUMI). (**E**-**G**) Two-dimensional UMAP representations of 13,942 single-cell transcriptomes generated by 10X Genomic RNAseq; the different panels show cells in function of their level of gene expression; selected genes are known to be specifically expressed in trophectoderm (**E**) including TFAP2C, KRT8, KRT18, CDX2, FABP3, and CLDN4, epiblast (**F**) including NANOG, FGF4, SOX15, GDF3, SFRP2, and SEMA6A, and primitive endoderm (**G**) including LAMA1, COL4A1, GATA4, FOXA2, SOX17, and GATA6.

#### Figure S2: Integration of rabbit, mouse and monkey single-cell RNAseq datasets

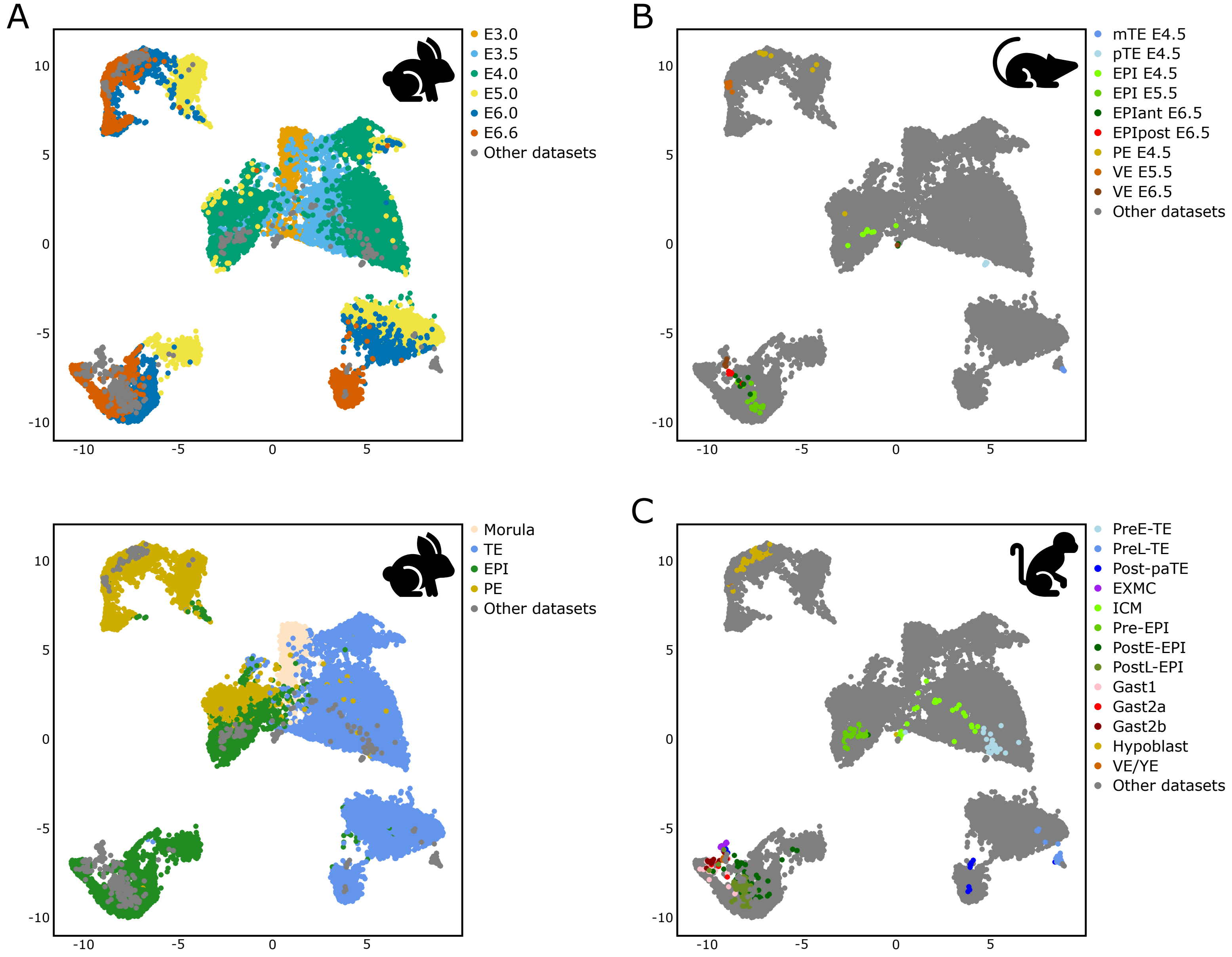


**Figure S2: Integration of rabbit, mouse and cynomolgus monkey single-cell RNAseq datasets.** UMAP representation of the integrated transcriptome dataset generated by merging the rabbit 10x dataset (our study), the mouse dataset (GSE63266) and the cynomolgus monkey dataset (GSE74767) (Nakamura et al., 2016). (**A**) Cells in our rabbit dataset are coloured according to either embryonic stages (upper panel) or embryonic lineages (lower panel). Cells from the mouse (**B**) or cynomolgus monkey (**C**) datasets are coloured according to their annotation in the published data. Cells from the other two datasets are shown in grey.

### Figure S3: Identification of late primitive endoderm and emergence of the germ line cells

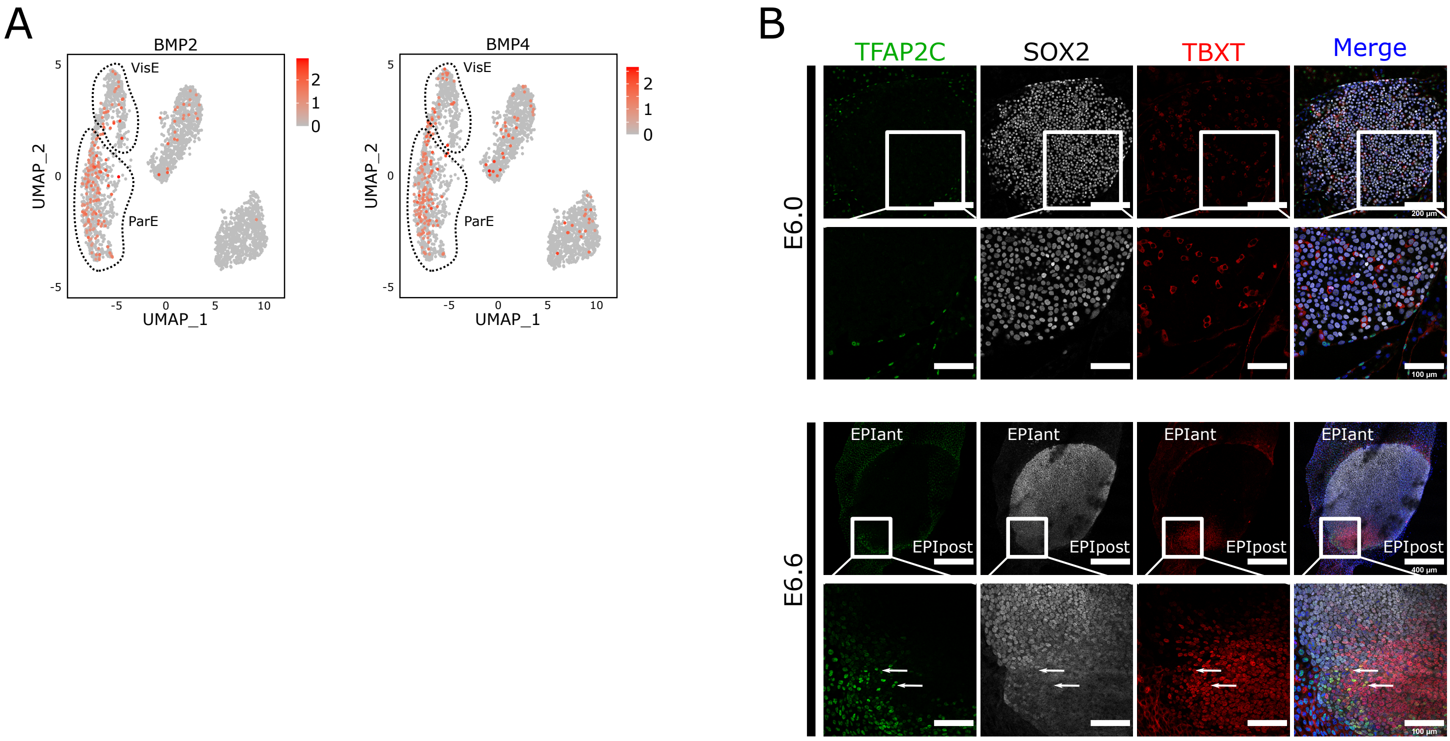


**Figure S3: Identification of late epiblast, late primitive endoderm and emergence of the germ line cells.** (**A**) UMAP representation of the PE cells transcriptome expressing either BMP2 or BMP4. VisE and ParE clusters are circled by dashed lines. (**B**) Immunofluorescence detection of SOX2, TFAP2C and TBXT in E6.0 (n = 5; scale bars: 400 µm, 100 µm, and 50 µm depending on enlargements) and E6.6 (n = 5; scale bars : 200 µm and 100 µm for enlargements) epiblasts. Arrows indicate TFAP2C/TBXT double-positive cells. Merge pictures include DAPI staining.

### Figure S4: Characterization of the pluripotency states.

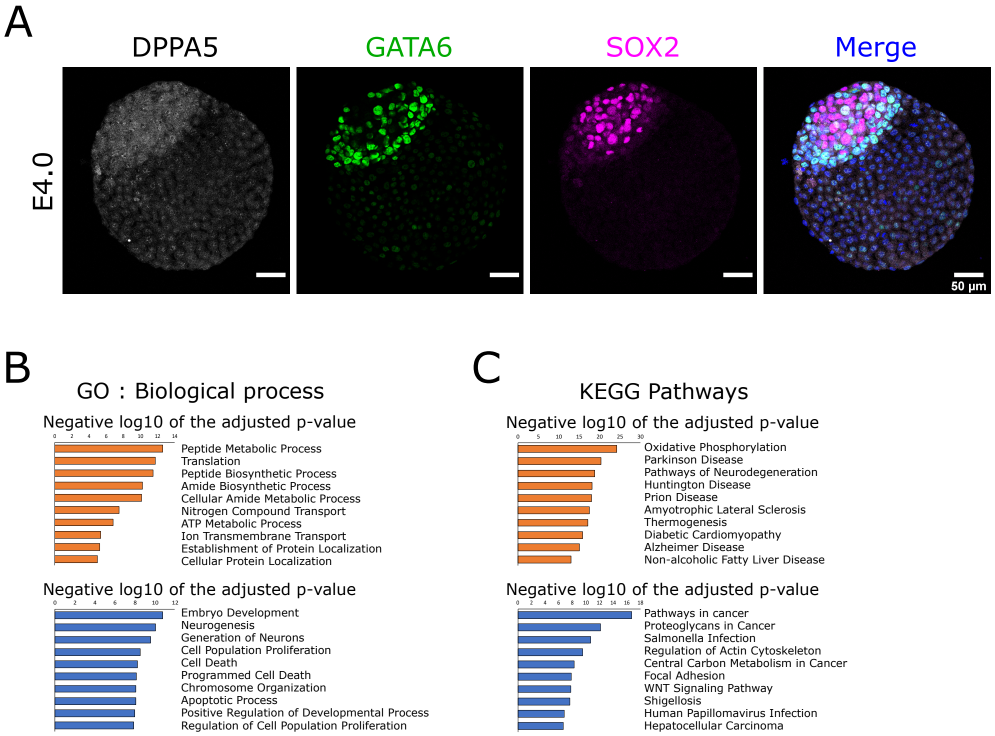


**Figure S4: Characterization of the pluripotency states.** (**A**) Immunofluorescence detection of DPPA5 (ICM-specific), GATA6 (PE-specific) and SOX2 (EPI-specific) in E4.0 blastocyst (n = 8). Merge image includes DAPI staining. Scale bars: 50 µm. (**B**) GO Biological Process enrichment analysis of the 500 most up- (red) and down-regulated (blue) genes in ICM and EPI_4.0 samples compared to EPI_ant, EPI_post and EPI_int samples in the 10x single-cell RNAseq dataset. (**C**) KEGG Pathways enrichment analysis of the 500 most up- (red) and down-regulated (blue) genes in in ICM and EPI_4.0 samples compared to EPI_ant, EPI_post and EPI_int samples in the 10x single-cell RNAseq dataset.

**Figure S5: Epigenetic and metabolic dynamics.**

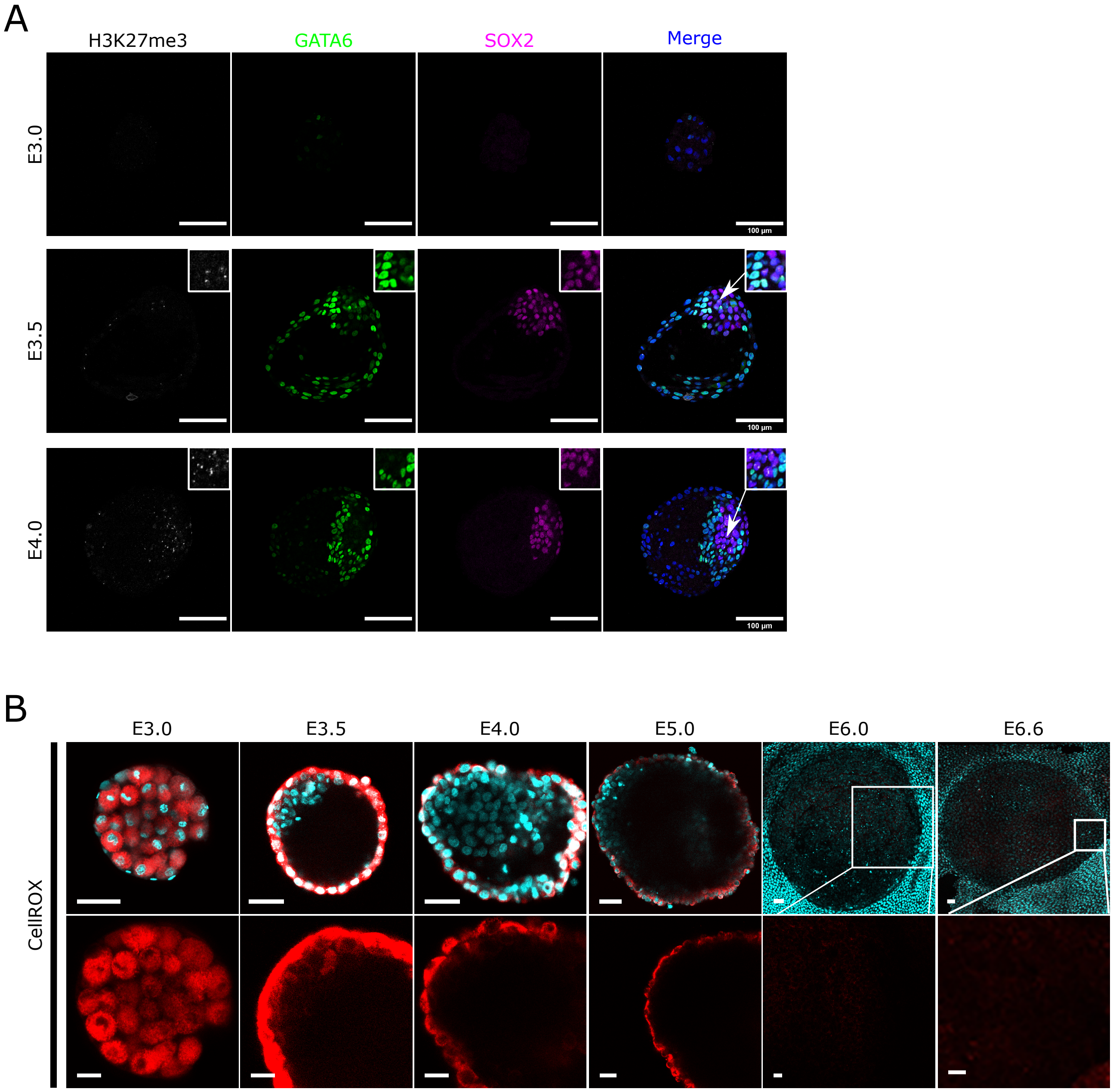


**Figure S5: Epigenetic and metabolic dynamics.** (**A**) Immunofluorescence detection of the histone 3 lysine 27 tri-methylation (H3K27me3) marks associated with X chromosome inactivation, GATA6 (PE-specific) and SOX2 (EPI-specific) in E3.0 (n = 9), E3.5 (n = 11) and E4.0 (n = 5) embryos. Merge pictures include DAPI staining. Scale bars: 100 µm. (**B**) Images of rabbit embryos (stages as indicated; n = 45, 5 to 10 per stage) treated with CellROX reagent to detect reactive oxygen species (ROS). Images in the top row (scale bars: 50 µm) show co-staining of ROS with CellROX reagent (red) and of nucleus with Hoechst (blue); Images in the bottom row (scale bars: 20 µm) show enlargement of ICM/epiblast cells after treatment with CellROX reagent only.

### Figure S6: Signaling pathways enrichment in EPI cells

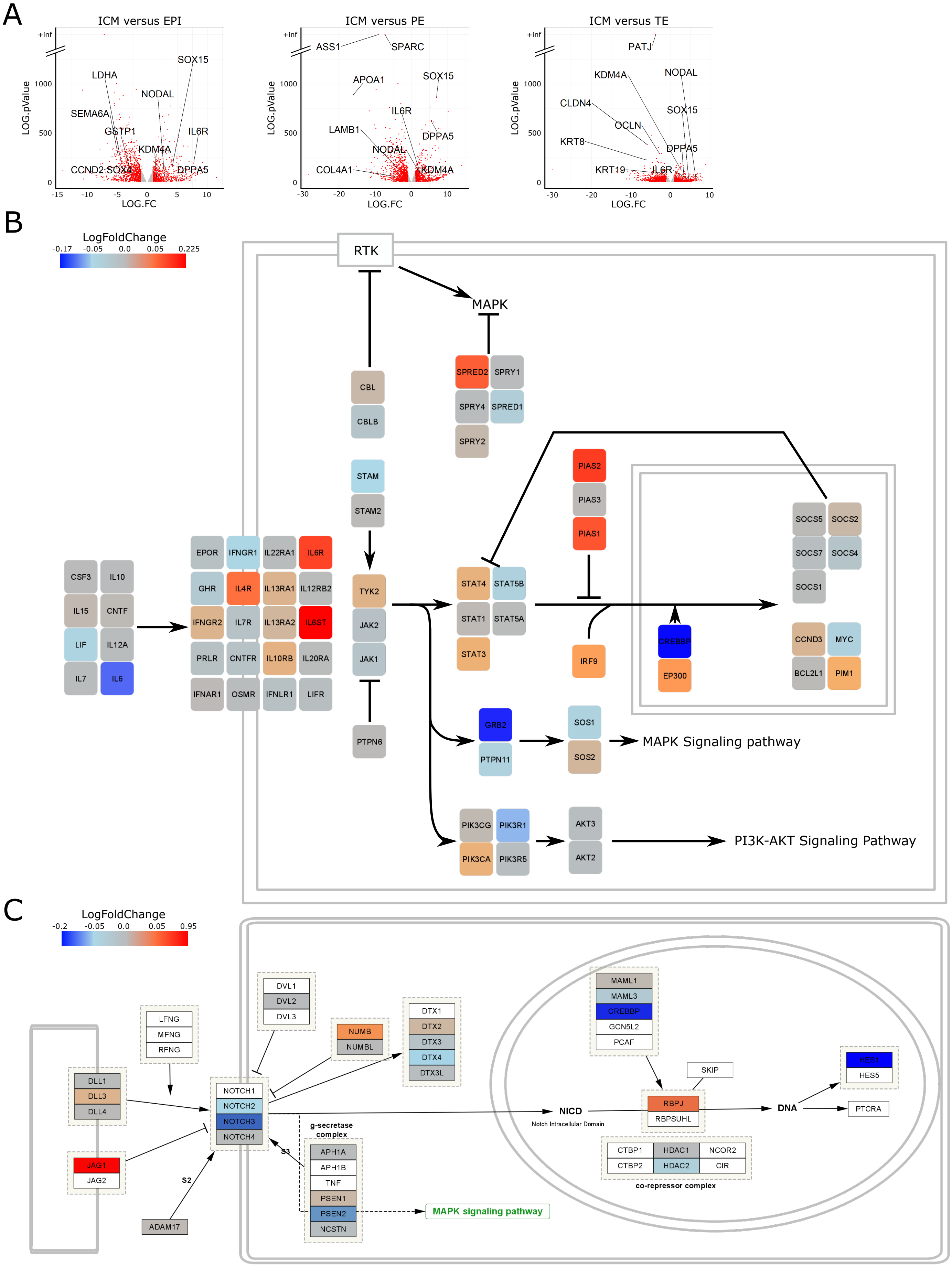


**Figure S6: Signalling pathways enrichment in EPI cells.** (**A**) Volcano plot representing the differential analysis between the ICM (E3.5/E4.0) and EPI (E6.0/E6.3/E6.6), PE (E6.0), or TE (E3.5/E4.0/E6.0/E6.6) in the bulk RNAseq dataset. The genes with a logarithm 10 fold change inferior to -1 or superior to 1 are coloured in red. (**B**,**C**) Schematic representation of JAK/STAT (**B**) and NOTCH (**C**) pathways highlighting differential expression (log Fold Change) of individual genes between ICM/EPI (E3.5/E4.0) and other cells (E3.5/E4.0/E5.0/E6.0/E6.6) in the 10X Genomics single-cell dataset.

### Figure S7: Expression of rabbit naïve pluripotency-specific genes in mouse and monkey embryos.

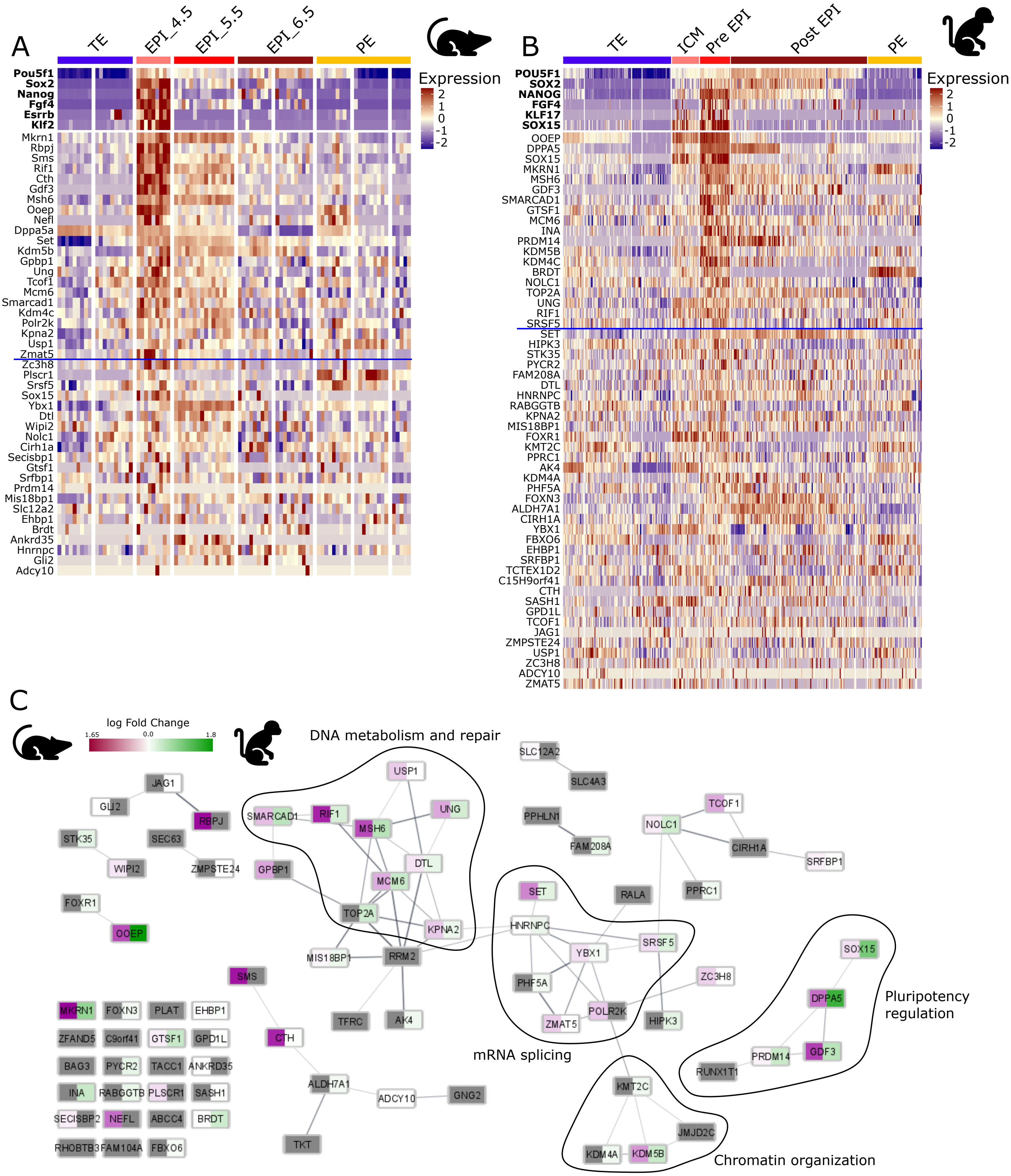


**Figure S7: Expression of rabbit naïve pluripotency-specific genes in mouse and monkey embryos.** (**A**) Heatmap of the 42 most differentially expressed genes (drawn from the list of 81 described in Fig. 8H-J), showing their transcript levels in mouse embryos (EPI cells at E4.5, E5.5 and E6.5, TE cells at E4.5, and PE cells at E4.5, E5.5 and E6.5; (Nakamura et al., 2016). (**B**) Heatmap representation of the 50 most differentially expressed genes (drawn from the list of 81 described in Fig. 8H-J), showing their transcript levels in cynomolgus macaque embryos (ICM cells at E6.0, pre-EPI cells at E7.0-E9.0, post_EPI cells at E13-E17, TE and PE at E7.0-E17; (Nakamura et al., 2016)). The blue line separates the significantly enriched genes (Fold change > 0.25) in the cynomolgus pre_EPI cluster and the mouse EPI_4.5 cluster. (**C**) STRING software network of the 81 rabbit naïve pluripotency-specific genes, with orthologs in cynomolgus macaque and mouse datasets (described in Fig. 8H-J), highlighting enrichment in the naïve embryonic cells (log Fold-Change) from mouse (purple) or cynomolgus macaque (green). In grey are genes that are not enriched in naïve embryonic cells of either species. Four main KEGG circuitries are shared by the three species analysed.
